# Constrained evolution of a core winter proteome across independently cold-adapted PACMAD grasses

**DOI:** 10.1101/2025.05.15.654294

**Authors:** Elad Oren, Jingjing Zhai, Travis E Rooney, Ruthie Angelovici, Charles O. Hale, Lara J. Brindisi, Sheng-Kai Hsu, Christine M. Gault, Jian Hua, Thuy La, Nicholas Lepak, Qin Fu, Edward S. Buckler, M. Cinta Romay

**Author notes:** **Corresponding authors**: Elad Oren, M. Cinta Romay. The author responsible for distribution of materials integral to the findings presented in this article is Elad Oren.

## Abstract

- Grasses in the PACMAD clade independently colonized cold environments from warm-climate ancestors, but whether their molecular responses to freezing reflect shared evolutionary solutions or lineage-specific innovations remains unknown. We used comparative proteomics to test whether protein-level cold responses show stronger cross-species conservation than previously observed at the transcript level.
- We quantified seasonal rhizome proteomes (winter vs summer) from five PACMAD species grown in a common garden exposed to sustained sub-zero temperatures, identified differentially abundant proteins, and compared fold-change magnitudes across species using orthogroup-based correlation analyses. We further examined LEA3 protein structure through hydropathy profiling and motif analysis.
- Shared cold-responsive proteins showed higher cross-species fold-change correlation (*ρ* = 0.80) than background proteins (*ρ* = 0.45), despite greater divergence in baseline abundance. LEA3 was the only ortholog elevated across all five species. Cold-tolerant species contained more tandem 11-mer repeats than the cold-sensitive maize, and two species accumulated multiple LEA3 paralogs, increasing total LEA3 abundance.
- Independent evolution of freezing tolerance in PACMAD grasses is governed by evolutionary constraints on protein-level response magnitude, reflecting the retention of an ancestral protective capacity. Structural divergence of LEA3 in maize suggests that transcriptional induction alone does not ensure freezing tolerance; functional protection likely requires intact motif architecture.

## Introduction

Maize (*Zea mays* ssp. mays), a vital crop for global food security, is susceptible to cold stress due to its tropical origins, limiting its yield in temperate regions. Early planting of maize can improve nitrogen capture, net primary productivity, and yield (Ojeda-Rivera et al., 2025). However, this strategy requires cold tolerance, which is lacking in current elite maize varieties (Lyons, 1973; Aroca et al., 2012). Cold and freezing tolerance has arisen independently multiple times within the Panicoideae, Aristidoideae, Chloridoideae, Micrairoideae, Aristavideae, and Danthonioideae (PACMAD) clade, a major lineage of predominantly C4 grasses (*Poaceae*) that diverged ∼64 Mya, with ancestral roots in the Afrotropics (Sage et al., 2011; Gallaher et al., 2022). These parallel origins of freezing tolerance provide a natural comparative framework. Among these, *Tripsacum*, the closest perennial and cold-tolerant relative of maize, having diverged about 650 Kya (Chen et al., 2022), is valuable for enhancing maize cold tolerance.

PACMAD grasses originated in warm-season climates unlike temperate diversified Pooideae and Danthonioideae (Kellogg, 2015). However, most transitions of PACMAD grasses to colder climates occurred in tropical mountains, where cold-adapted lineages converged with temperate migrants (Schubert et al., 2020). Each PACMAD genus exhibits distinct evolutionary trajectories, polyploid histories, and biogeographic origins, having independently evolved robust cold and freezing tolerance over the past 20-30 million years (Fig. 1e; Gallaher et al., 2022; Stitzer et al., 2025).

**Fig. 1.**
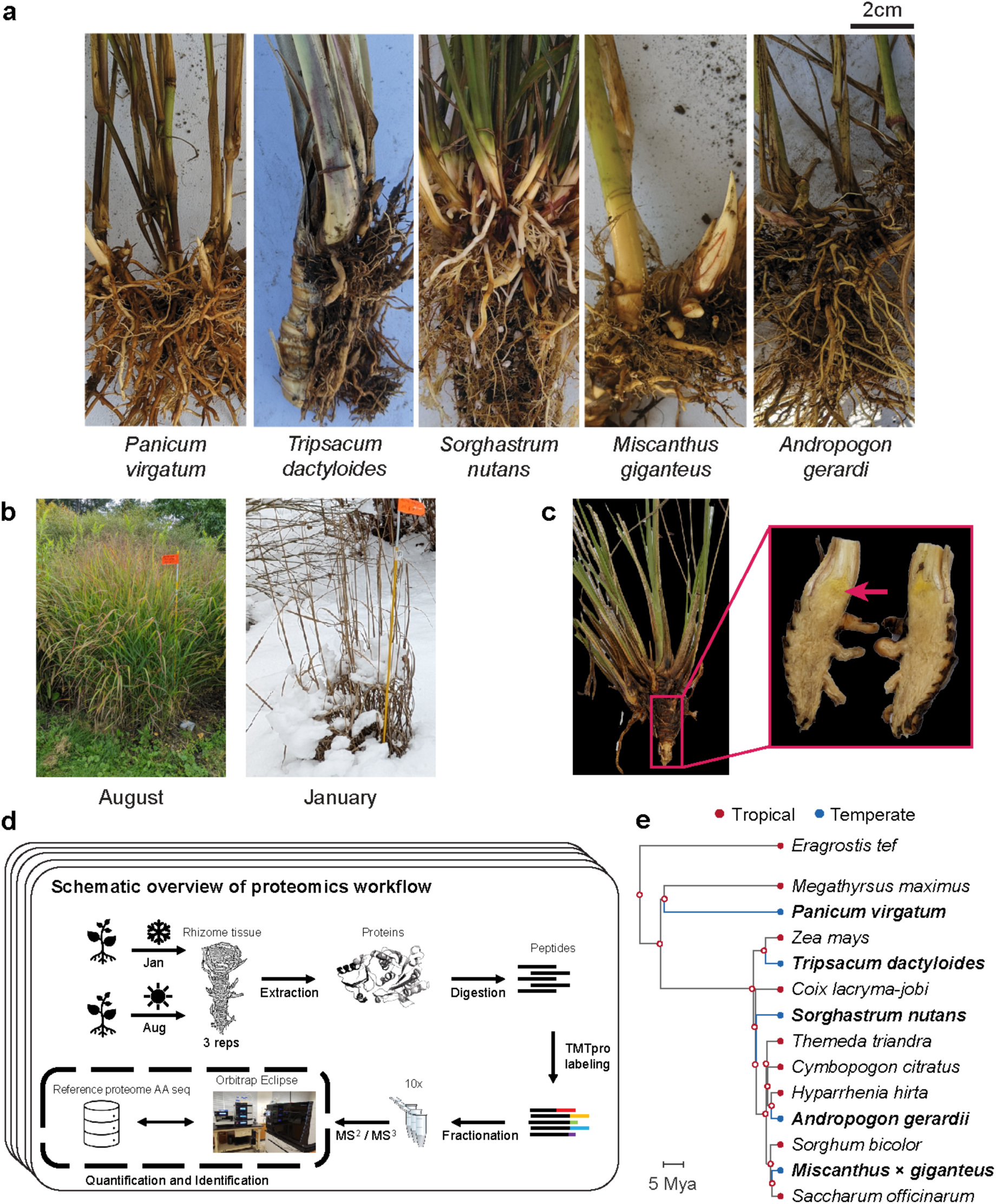
Overview of sampling and proteomics workflow. **(a)** Rhizomes from the five species sampled in the study: *Tripsacum dactyloides* and *T. floridanum* hybrids, *Panicum virgatum*, *Andropogon gerardi*, *Miscanthus × giganteus*, and *Sorghastrum nutans*. **(b)** Field conditions during the two sampling seasons, illustrating the active growth in August and dormancy in January. **(c)** Example of the rhizome tissue sampled for proteomics, shown here for *Tripsacum*. The red arrow highlights the specific region collected. **(d)** Schematic overview of the proteomics workflow. Rhizome samples were collected each season, followed by protein extraction, digestion, and TMTpro labeling. Peptides were fractionated and analyzed via RTS-SPS-MS^3^ using an Orbitrap Eclipse mass spectrometer. Each species’ reference proteome was used for peptide and protein assignment, enabling quantification and identification. **(e)** Phylogenetic relationships among the five study species (bold) and tropical relatives, rooted on *Eragrostis tef* (Chloridoideae). Red, tropical; Blue, temperate and freezing tolerant.

Plant exposure to cold stress, including chilling (>0°C) and freezing (<0°C) temperatures, triggers physiological and molecular responses. Chilling stress disrupts membrane fluidity, impairs enzyme activity, and inhibits photosynthesis (Aroca et al., 2012), whereas freezing stress induces ice crystal formation in extracellular spaces, leading to cellular dehydration, membrane damage, and ultimately, cell death (Pearce, 2001). Many temperate plants undergo cold acclimation, resulting in adaptations such as membrane lipid modifications (Degenkolbe et al., 2012; Barnes et al., 2016), accumulation of compatible solutes (Koster & Lynch, 1992), enhanced antioxidant enzyme activity (Suzuki & Mittler, 2006) and protective protein accumulation (Puhakainen et al., 2004; Xu et al., 2022). Plant cold perception involves plasma membrane modifications, including fluidity changes, Ca²⁺ channel activation, and cytosolic acidification, driving lipid and transcriptomic responses (Mori et al., 2018; Barnes et al., 2023). A widely conserved mechanism underpinning freezing tolerance is the activation of the C-repeat binding factor/dehydration-responsive element-binding (CBF/DREB) signaling pathway, which induces expression of cold-responsive (COR) genes (Thomashow, 1999; Nakashima & Yamaguchi-Shinozaki, 2006).

Studies in PACMAD grasses reveal both broadly conserved and species-specific molecular adaptations associated with this pathway. *Panicum virgatum* utilizes a zinc finger transcription factor, PvC3H72, to enhance ICE1-CBF-COR signaling, significantly increasing cold resilience (Xie et al., 2019). Additionally, calcium signaling pathways involving calmodulin and MEKK1 provide early response to cold stress (Ranaweera et al., 2022; Wu et al., 2024). *Pennisetum giganteum* maintains membrane fluidity at low temperature through the accumulation of unsaturated fatty acids, such as linoleic and α-linolenic acids (Zhang et al., 2017; Li et al., 2020). Protective proteins, such as late embryogenesis abundant (LEA) proteins, dehydrins, and antioxidant enzymes, are accumulated in response to freezing conditions, reducing oxidative injury and cellular dehydration (Liu et al., 2016; Xie et al., 2019). Species-specific adaptations include enhanced proline biosynthesis via the ornithine pathway in *Sorghum bicolor* (Vera-Hernández et al., 2018) and elevated activity of phenylpropanoid pathway enzymes (PAL and CAD) in *Miscanthus* to protect cells during freezing stress (Domon et al., 2013).

Previous studies in PACMAD grasses have focused mainly on transcriptomic and metabolomic adaptations. However, proteomic responses remain largely unexplored. As the functional products of genes, proteins integrate transcriptional, translational, and post-translational regulation. Given the persistent nature of cold stress, proteomic analysis uniquely captures the accumulation of proteins reflecting acclimation and sustained adaptive responses. Furthermore, Cope et al. (2025) demonstrated that proteins face stronger evolutionary constraints than transcripts, with protein abundances being highly conserved across species while mRNA levels can vary without substantially affecting protein output. This suggests that shared cold adaptive responses may be more readily detected at the protein level than at the transcript level, making proteomics essential for understanding adaptive processes such as those underlying cold or freezing tolerance. Our multi-species study examines rhizome protein accumulation across seasonal variation to identify conserved freezing-tolerance mechanisms in PACMAD grasses, testing whether freezing tolerance across these lineages reflects retention of shared ancestral molecular responses or lineage-specific innovations.

## Materials and Methods

To investigate proteomic responses to seasonal changes across PACMAD grasses, we conducted a comparative analysis of rhizomes from five species under winter (January, with lows reaching –29°C) and summer (August, with highs of 26°C) conditions. The species analyzed (Fig. 1a) were interspecific and intraspecific hybrids of *Tripsacum dactyloides* (L.) L. and *Tripsacum floridanum* Porter ex Vasey, *Andropogon gerardi* Vitman (cv. ‘Sentinel’), *Miscanthus* × *giganteus* J.M.Greef & Deuter ex Hodk. & Renvoize, *Panicum virgatum* L. (cv. ‘Dust Devil’), and *Sorghastrum nutans* (L.) Nash (cv. ‘Golden Sunset’).

### Phylogenetic context of freezing tolerance

To illustrate independent origins of freezing tolerance in each of the five study species rather than from a single common ancestor, we constructed phylogenetic context by pruning the 1,153-tip ASTRAL-Pro nuclear species tree (Arthan et al., 2024) to 14 representative species (Fig. 1e): Paniceae (2 species) and Andropogoneae (11 species from 7 subtribes) and *Eragrostis tef* (Chloridoideae) as outgroup. Of these 14, only five temperate species were sampled for proteomics analysis (*P. virgatum*, *T. dactyloides*, *S. nutans*, *A. gerardi*, and *M. giganteus*); the remaining nine tropical relatives (*Megathyrsus maximus*, *Zea mays*, *Coix lacryma-jobi*, *Themeda triandra*, *Cymbopogon citratus*, *Hyparrhenia hirta*, *Sorghum bicolor*, *Saccharum officinarum*) were included to demonstrate that each study species sits on a separate branch from its nearest warm-climate relative, confirming five independent transitions to freezing tolerance. *Sorghastrum fuscescens* served as a positional proxy for *S. nutans*, which was absent from the tree, and *M. giganteus’s* position was determined by its maternal progenitor, *Miscanthus sinensis*. Thermal ecology was classified as tropical or temperate based on native range descriptions in Kellogg (2015).

### Plant material and environmental conditions

Plants were grown outdoors at two field sites in Ithaca, New York (USDA Hardiness Zone 6a), where they experienced natural seasonal fluctuations, with an average winter temperature of – 5°C (lows reaching –29°C) and summer temperatures averaging 20°C (highs of 26°C) over the five years preceding sampling (Northeast Regional Climate Center, 2025). *Tripsacum* hybrids were cultivated at Cornell University Caldwell Field, while *A. gerardi, M. giganteus, P. virgatum*, and *S. nutan*s were grown at the Cornell Botanic Gardens. Environmental data were obtained from the Northeast Regional Climate Center (Station USC00304174) and the New York State Mesonet (Station GROT; Brotzge et al., 2020).

### Sampling and experimental design

Rhizome tissue was collected during two sampling campaigns in 2019-2020 and 2022, timed to capture distinct physiological states: winter dormancy and summer active growth. Summer samples were collected in August 2019 and August 2022 during active growth; winter samples in January 2020 and December 2022 during dormancy. Soil temperatures and precipitation data for each sampling date are reported in Supporting Information Methods S1.

At summer sampling, plants displayed active vegetative rhizome growth with no visible drought stress; winter samples were dormant with hardened exteriors (Fig. S1). Our study was conducted in two phases. Initial sampling of *Tripsacum* hybrids in 2019 provided evidence of conserved winter protein accumulation, prompting expansion in 2022 to both replicated *Tripsacum* sampling and four additional PACMAD species with independent evolutionary histories of cold adaptation.

Rhizome tissue (minimum 5 g fresh weight) was collected from the internal mid-sections of rhizomes using sterile pruning scissors. Sampling was standardized to exclude apical tips, fine roots, and necrotic zones to ensure reproducibility and tissue comparability across species (Fig. 1c and Fig. S1). Samples were immediately placed in pre-labeled 15 ml tubes, transported on dry ice, and stored at −80°C within 4 hours of collection. Biological replicates were defined as tissue collected from independent clonal plants. Sample sizes and genotypes are detailed in Table 1. For *A. gerardi* (’Sentinel’), *M. giganteus*, and *P. virgatum* (’Dust Devil’), three biological replicates were sampled per season from established accessions at Cornell Botanic Gardens. *S. nutans* (’Golden Sunset’) was represented by two biological replicates per season due to winter mortality of one clonal plant.

**Table 1.** Summary of rhizome proteomics datasets for *Tripsacum* hybrids, *Andropogon gerardi*, *Miscanthus* × *giganteus*, *Panicum virgatum*, and *Sorghastrum nutans*, including number of samples per season, total proteins quantified, proteins shared across seasons, and differentially accumulated proteins (DAPs) accumulated (up) or depleted (down) in winter.

| Species | Year | N samples | Genotype / Cultivar | Entry code | Clonal reps | Proteins quantified |  | DAP up | DAP down |
| --- | --- | --- | --- | --- | --- | --- | --- | --- | --- |
|  |  |  |  |  |  | Total | Shared winter/summer |  |  |
| *Tripsacum* spp. | 2019 | 8 | *T. dactyloides* 'IA_Pete_2' × *T. floridanum* | C002 | 1 | 4,451 | 4,451 | 123 | 175 |
|  |  |  | *T. dactyloides* 'KS_C9_9' × *T. floridanum* | C004 | 1 |  |  |  |  |
|  |  |  | *T. dactyloides* 'KS_B6_1' × *T. floridanum* | C022 | 3 |  |  |  |  |
|  |  |  | *T. dactyloides* 'KS_C9_10' × *T. floridanum* | C031 | 1 |  |  |  |  |
|  |  |  | *T. dactyloides* 'IA_Pete_1' × *T. dactyloides* 'FL' | C071 | 1 |  |  |  |  |
|  |  |  | *T. dactyloides* 'TX_HC_3' × *T. dactyloides* 'MO_TP' | C076 | 1 |  |  |  |  |
| *Tripsacum* spp. | 2022 | 24 | *T. dactyloides* 'IA_Pete_2' × *T. floridanum* | C002 | 3 | 3,468 | 3,054 | 100 | 141 |
|  |  |  | *T. dactyloides* 'KS_C9_9' × *T. floridanum* | C004 | 3 |  |  |  |  |
|  |  |  | *T. dactyloides* 'KS_B6_1' × *T. floridanum* | C022 | 3 |  |  |  |  |
|  |  |  | *T. dactyloides* 'KS_C7_3' × *T. floridanum* | C027 | 3 |  |  |  |  |
|  |  |  | *T. dactyloides* 'IL_10_1' × *T. floridanum* | C043 | 3 |  |  |  |  |
|  |  |  | *T. dactyloides* 'IL_14' × *T. dactyloides* 'FL' | C065 | 3 |  |  |  |  |
|  |  |  | *T. dactyloides* 'IA_Pete_1' × *T. dactyloides* 'FL' | C071 | 3 |  |  |  |  |
|  |  |  | *T. dactyloides* 'TX_HC_3' × *T. dactyloides* 'MO_TP' | C076 | 3 |  |  |  |  |
| *Andropogon gerardi* | 2022 | 3 | Sentinel | 01-266*A | 3 | 3,734 | 3,733 | 109 | 76 |
| *Miscanthus* × *giganteus* | 2022 | 3 | - | 88-363*A | 3 | 2,372 | 2,273 | 58 | 83 |
| *Panicum virgatum* | 2022 | 3 | Dust Devil | 01-277*B | 3 | 4,071 | 4,068 | 174 | 218 |
| *Sorghastrum nutans* | 2022 | 2 | Golden Sunset | 22-095*A | 2 | 2,641 | 2,464 | 66 | 56 |

### Tripsacum hybrid population

*Tripsacum* rhizomes were sampled from seven-year-old F₁ hybrid progenies that had survived multiple freezing winters at Cornell University’s Caldwell Field. The hybrids were derived from controlled crosses between northern founders collected from Iowa, Kansas, Illinois, and Missouri, where winters experience heavy frost, and southern founders collected from Florida and Texas, with subtropical climates. Most crosses were interspecific, between *T. dactyloides* subsp. *dactyloides* (northern) and *T. floridanum* (southern); additional crosses were intraspecific, between northern and southern populations of *T. dactyloides* (Table 1). In 2019, we sampled eight individuals per season representing six distinct hybrid genotypes from five cross combinations, prioritizing coverage of genetic diversity across the hybrid population. In 2022, we sampled 24 individuals per season comprising eight hybrid genotypes with three clonal replicates each, balancing genetic diversity with robust statistical replication.

### F₂ Seedlings for RNA sequencing

F₂ seeds used for seedling freezing tolerance screening and RNA sequencing were generated from the same *Tripsacum* hybrids described above. Due to the limited seed set achieved through self-pollination (∼500 seeds over the entire season), F₁ plants were allowed to open-pollinate. Seeds were harvested from individual maternal plants, germinated in trays, and grouped into F₂ families based on maternal lineage. Six-week-old F₂ seedlings sharing the same maternal F₁ parent were selected as experimental material for freezing assays and RNA extraction.

### Proteomics data

Proteomic profiling was conducted using tandem mass tag (TMTpro) labeling and Orbitrap Eclipse Real Time Search (RTS)-SPS-MS^3^ technology. The experimental design included samples collected in both winter (January) and summer (August), with each species represented by at least three biological replicates with a larger number of genotypes analyzed for *Tripsacum* across two years (see Table 1 and Fig. 1d).

### Protein Extraction and Digestion

Fresh rhizome tissue was ground in liquid nitrogen and subjected to phenol-based protein extraction with protease and phosphatase inhibitors, followed by ammonium acetate/methanol precipitation. Protein pellets were dissolved in 8 M urea, quantified by BCA assay, and digested using the S-Trap protocol with trypsin. Digested peptide mixtures were labeled using TMTpro reagents: TMTpro 16-plex for 2019 Tripsacum samples and TMTpro 18-plex for 2022 samples. The 2022 samples were processed across seven separate TMTpro 18-plex batches: four batches for Tripsacum and three batches for the remaining four species, with winter and summer samples from each species always included within the same batch. Detailed buffer compositions, wash steps, digestion parameters, and labeling procedures are provided in Supporting Information Methods S1.

### LC-MS/MS analysis and database searching

Labeled peptides were fractionated by high-pH reversed-phase chromatography (48 fractions concatenated into 10 pools) and analyzed by nanoLC coupled to an Orbitrap Eclipse mass spectrometer using the RTS-SPS-MS³ method (Fu et al., 2021). Raw data were processed in Proteome Discoverer 2.5 using the Sequest HT search engine against species-specific reference proteomes (see Peptide Mapping and Reference Proteome Assignment). Peptide spectral matches were filtered at 1% FDR (Percolator), and protein identifications required a minimum of two distinct peptides. Protein quantification was based on summed reporter ion intensities of unique and razor peptides. Full search parameters, modification lists, mass tolerances, and reporter ion filtering criteria are detailed in Methods S1.

Given the inherent biological variability of a multi-species field study, we applied a stringent threshold for differential abundance. Differentially abundant proteins (DAPs) were determined based on |log₂ fold change| ≥ 1 and adjusted P-value < 0.05, to prioritize high-confidence candidates with robust seasonal responses over minor fluctuations that may reflect transient environmental noise rather than conserved adaptive strategies. This approach helps address the statistical power imbalance between the deeply sampled Tripsacum dataset (n≈32) and the other grass lineages (n=3). While the higher power in Tripsacum inherently yields a larger list of statistically significant changes, our comparative analysis defines conservation based on the intersection of species. Consequently, the identification of conserved proteins is constrained by the more limited statistical power of the smaller datasets, suggesting that the core response network we report represents a high-confidence set of shared adaptations rather than an exhaustive catalog.

### Peptide Mapping and Reference Proteome Assignment

Initial Sequest HT searches were performed against species-specific reference proteomes (see above). For species pooled within the same TMT batch due to <15% predicted proteome divergence, such as *A. gerardi* and *M. giganteus*, spectra were searched against the *A. gerardi* proteome. To improve protein recovery for *M. giganteus*, peptides were re-mapped to the *M. sinensis* proteome using DIAMOND BLASTp in “ultrasensitive” mode. Proteins were reassigned to their best-matching *M. sinensis* counterparts based on aggregated peptide evidence (see Supporting Information Code S1–S3; GitHub repository).

All final analyses were conducted using species-specific reference proteomes: *Andropogon gerardi* (Ag-CAM1351-DRAFT-PanAnd-1.0), *Miscanthus × giganteus* (Msinensis_497_v7.0), *Panicum virgatum* (Pvirgatumvar_WBCHAP1_778_v1.0), *Sorghastrum nutans* (Sn-CAM1369-DRAFT-PanAnd-1.0), and *Tripsacum dactyloides* (Td-FL_9056069_6-DRAFT-PanAnd-1.0); genome assemblies and annotations are listed in the Data Availability section.

### Protein abundance normalization and statistical analysis

Protein intensities were normalized using the “total peptide amount” method in Proteome Discoverer 2.5. Differential abundance between winter and summer was assessed using one-way ANOVA followed by Tukey’s HSD post hoc test, with multiple testing correction by the Benjamini-Hochberg procedure. For the multi-batch *Tripsacum* 2022 dataset, statistical testing was performed within each batch and P-values were combined using Fisher’s method; only proteins quantified in at least 3 of 4 batches were retained. Proteins quantified in both seasons within each species were considered differentially abundant at |log₂FC| ≥ 1 and adjusted *P* < 0.05.

PCA was performed on log₂-transformed protein abundances (Table S1) and on orthogroup-level data detected in >50% of samples per species (Fig. S2). For Tripsacum, batch effects were removed via residuals from a linear model prior to PCA; because season was the dominant factor (Fig. S2b), all Tripsacum samples were pooled across years. For cross-species comparison, orthogroups were assigned using OrthoFinder (Emms & Kelly, 2019), with fold-change values averaged when multiple proteins mapped to the same orthogroup. All analyses were performed in R v4.4.3.

### Cross-species correlation analysis

To assess evolutionary constraint on cold-responsive proteins, we compared cross-species conservation of seasonal response magnitude between shared cold-responsive orthogroups and background proteins, following the framework of Cope et al. (2025).

For each orthogroup quantified in at least four of five species, we calculated pairwise Spearman correlations of log₂ fold-change (winter/summer) across all species pairs. Orthogroups were classified as “shared DAPs” if differentially abundant (|log₂FC| ≥ 1, adjusted *P* < 0.05) in two or more species, or as “background” if differentially abundant in at most one species. Distributions of pairwise correlations were compared between shared DAPs and background using the Wilcoxon rank-sum test. The same approach was applied to basal protein abundance (summer and winter separately) to distinguish constraint on response dynamics from constraint on absolute abundance levels.

### Functional annotation and GO enrichment

Orthogroups were assigned using OrthoFinder (v2.5.5; Emms & Kelly, 2019), with DIAMOND (v2.1.9; Buchfink et al., 2021) for all-vs-all sequence comparisons across all five species. Orthogroups served as the basis for cross-species comparisons and functional propagation of annotations. DAP sequences were annotated using DIAMOND BLASTp against the SwissProt with gene ontology (GO) terms extracted from UniProt. For DAPs lacking SwissProt hits, functional predictions were generated using PANNZER2 (Törönen & Holm, 2022), run independently for each species. Annotation of non-significant proteins is detailed in Methods S1.

GO term enrichment analysis was performed using the topGO R package (v2.50.0; Alexa & Rahnenfuhrer, 2017), with the “elim” algorithm with Fisher’s exact test, separately for winter-elevated and winter-reduced proteins against all quantified proteins. Terms with a minimum node size of 10 and a P-value < 0.01 were considered significant. Due to the hierarchical nature of topGO’s algorithm, multiple test correction was not necessary (Alexa & Rahnenfuhrer, 2017).

### Candidate protein exploration methods

Barnes et al. (2023) showed that cold-responsive transcriptional and proteomic changes often overlap with broader stress mechanisms involving damage, osmotic imbalance, and acidification. To improve biological resolution, we curated DAPs into nine functional categories based on known roles in freezing tolerance (Orvar et al., 2000; Iwaya-Inoue et al., 2018; Schubert et al., 2020): (1) protein aggregation and membrane stability, (2) protein folding and chaperone activity, (3) metabolism and osmoregulation, (4) detoxification and ROS scavenging, (5) lipid metabolism, (6) antifreeze and cell wall modification, (7) cold signal transduction, (8) cytoskeletal organization, and (9) other/uncategorized. Assignments were informed by protein descriptions, GO terms, PANNZER2 predictions, and domain feature review (Fig. S3).

### Selecting proteins for traits on trees

Proteins for phylogenetic trait mapping were identified by text-mining PANNZER2 annotations for GO terms associated with LEA proteins, HSPs, and EF1. Full-length sequences were extracted from reference proteomes using seqkit grep (v2.0), aligned with MAFFT (v7.475), and used to infer maximum-likelihood trees with raxmlHPC (RAxML-8.2.13, `PROTGAMMAJTT`, 100 bootstraps). Trees were rooted on appropriate outgroups and visualized with ggtree, mapping log₂FC values as tip-point colors. Alignment and tree-building parameters are detailed in Methods S1.

### RNA-seq processing and differential expression analysis

Transcriptome data for maize chilling responses across tissues (leaf, root, top-crown, and bottom-crown) were derived from Xue et al. (2021) using 3′ RNA-seq (5C vs. 22C). For *Tripsacum*, we analyzed leaf transcriptome data from a segregating freezing-tolerant bulk of F₂ seedlings derived from the same *Tripsacum* hybrids used in our rhizome analysis. Plants from seven F₂ families were bulked as freezing-tolerant or intolerant based on survival after exposure to –8°C for three hours. Within the freezing-tolerant bulk only, RNA was collected from 6-week-old seedlings at control temperature (22°C) and after 7 days of cold acclimation (5°C), with expression compared between these conditions (tol_5C vs. tol_22C). The second transcriptome from *Tripsacum* ‘Pete’ (leaf, root, top-crown, and bottom-crown) was generated using the same methodology (5°C vs. 22°C) as in Xue et al. (2021) and is currently unpublished.

F₂ seedling leaf RNA was extracted following Gault et al. (2018). Paired-end reads were quality-trimmed using Trimmomatic (v0.32; Bolger et al., 2014), aligned to the T. dactyloides genome (Td-FL_9056069_6-DRAFT-PanAnd-1.0) using STAR v2.7.10b, and quantified with featureCounts v2.0.3. Differential expression was assessed using DESeq2 (Love et al., 2014), modeling temperature and genotype as main effects. Genes were considered differentially expressed at adjusted P < 0.05 (Benjamini-Hochberg) and |log₂FC| > 1. Trimming parameters, STAR alignment settings, and the full DESeq2 model are detailed in Methods S1.

### Structural and Biophysical Analysis of LEA3 Proteins

Because LEA3 emerged as the only differentially abundant protein consistently elevated across all five species (see Results) we performed targeted structural and biophysical analyses to characterize its repeat architecture and cryoprotective potential. LEA3 orthologs from all five study species plus *Zea mays* were aligned using MAFFT v7.520 with default settings.

Predicted three-dimensional structures for the *Tripsacum dactyloides* (Td00001aa022594_T002) and *Zea mays* (Zm00001eb294480) orthologs were generated using AlphaFold3 (Abramson et al., 2024) with default parameters and full length amino acid sequence as input. Hydrophobicity was visualized in PyMOL (v2.5) using a custom script based on the Eisenberg scale, which colored residues according to side-chain properties. Hydropathy plots were generated using both ProtScale (ExPASy) and AmphipaSeek (Sapay et al., 2006), with a sliding window size of 11 amino acids corresponding to the LEA3 motif length. Hydrophobic peaks associated with conserved LEA motifs were annotated across both sequence alignments and structural models.

## Results

To examine seasonal proteomic responses in PACMAD grasses, we analyzed rhizome tissues of five species using a proteomic shotgun approach during the winter (January) and summer (August) of 2019-2020 and 2022 (Fig. 1) in Ithaca, New York (USDA Hardiness Zone 6a). The exposure to deep-freezing winters ensured that the proteomic responses captured were reflective of natural seasonal variation (Fig. 1b).

Protein abundance analysis was performed in batches (see Methods). Across all five species a total of 20,737 proteins were quantified, but for differential accumulation across seasons we included only those quantified in both winter and summer samples (Table 1, Fig. S4). The number of identified proteins ranged from 2,372 to 4,451 per species. Differentially abundant proteins (DAPs) varied across species, with 58–174 proteins significantly increased and 56–218 decreased in abundance under freezing conditions (Table 1). PCA of 745 shared orthogroups confirmed clustering by species and season (Fig. S2); batch-corrected *Tripsacum* samples (1,889 proteins) clustered by season regardless of hybrid background (Fig. S2b). Although fewer proteins were identified in *Sorghastrum nutans*, reflecting its draft genome status (Table S2), robust seasonal signatures were detectable across all five lineages (Fig. S2a).

### Cross-Species comparative proteomic analysis reveals conserved seasonal protein accumulation patterns

We restricted our comparative analysis to the 18,454 proteins assigned to 4,888 orthogroups identified across species (Tables S3 and S4). Seasonal fold-changes were positively correlated across all species pairs (*ρ* = 0.37-0.69; Fig. 2a), with specific orthogroups—including LEA3, GST, HSP1, and PEBP—consistently identified as differentially abundant. To test whether this conservation reflects evolutionary constraint on cold response, we compared fold-change correlations between shared DAPs (differential abundance in ≥2 species; n = 135 orthogroups) and background proteins (≤1 species; n = 1,548). Shared DAPs showed substantially higher correlation in seasonal response magnitude (*ρ* = 0.80) than background (*ρ* = 0.45; *P* = 1.1×10⁻⁵; Fig. 2b). Notably, this constraint did not extend to basal abundance, where shared DAPs showed slightly lower correlation than background (*ρ* = 0.62 vs 0.68; *P* = 0.004). This pattern—tight constraint on response dynamics but flexibility in baseline levels—suggests that independent cold adaptations are governed by conserved regulation of a core set of protective proteins. This result was robust to the exclusion of *S. nutans*, which had fewer biological replicates (n = 2 per season; *P* = 0.002).

**Fig. 2.**
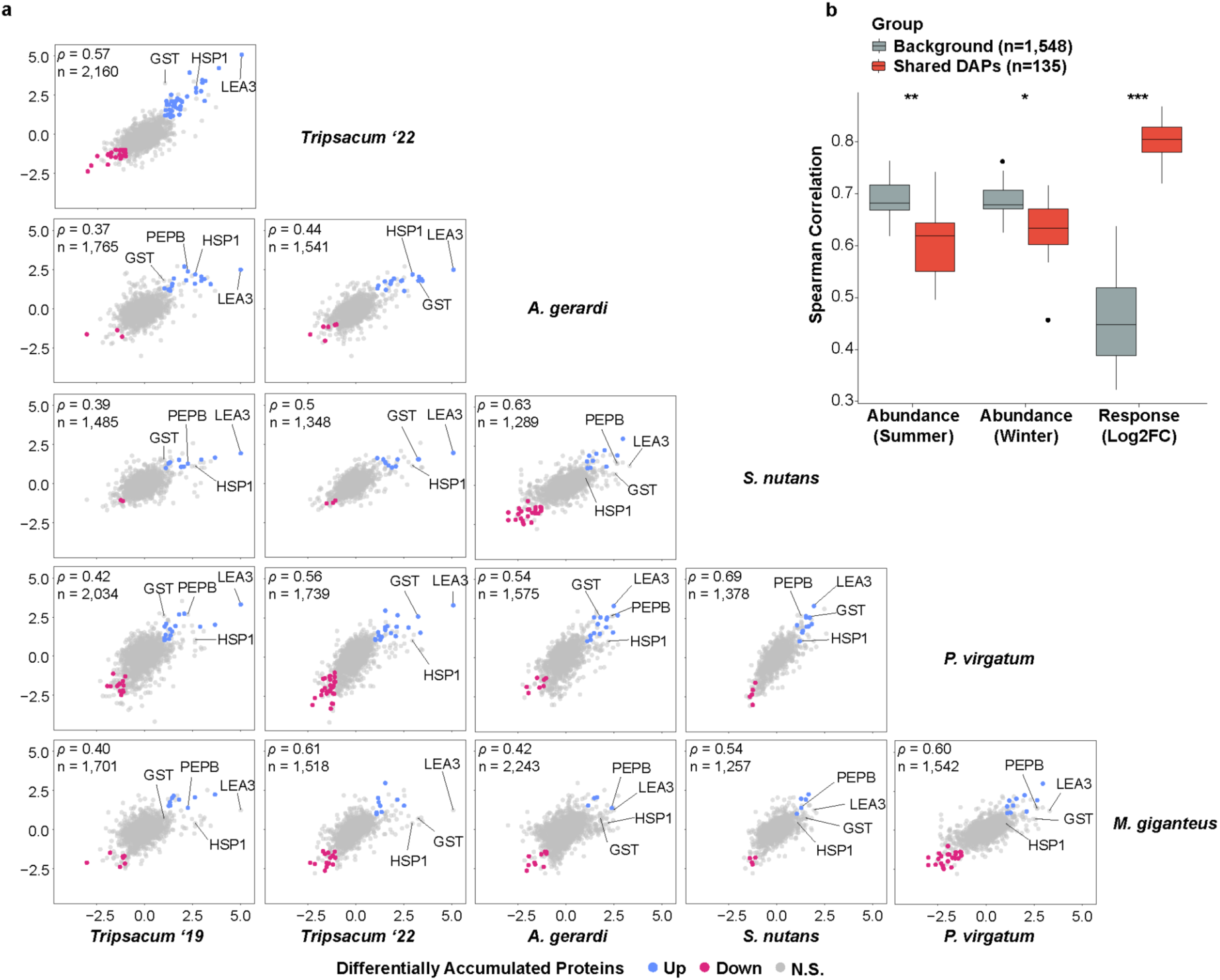
Conservation of seasonal protein accumulation in rhizomes of *Tripsacum dactyloides* and *T. floridanum* hybrids, *Andropogon gerardi*, *Miscanthus* × *giganteus*, *Panicum virgatum*, and *Sorghastrum nutans*. (a) Pairwise comparisons of log_2_ fold-change (log₂FC winter/summer) in differentially accumulated orthogroups across species. Scatterplots display the correlation of orthogroup-level log_2_FC values between species, with shared DAPs elevated in winter shown in blue, reduced in red, and non-significant proteins in gray. Each panel represents a comparison between two species, with the number of shared orthogroups (*n*) and Spearman’s correlation coefficient (ρ) displayed. **(b)** Pairwise Spearman correlations for orthogroups quantified in at least four of five species, comparing shared DAPs (red; differentially abundant in ≥2 species) versus background orthogroups (gray; differentially abundant in ≤1 species). Correlations are shown for summer abundance, winter abundance, and seasonal response magnitude (log₂FC winter/summer). Wilcoxon rank-sum test: *P < 0.05, **P < 0.01, ***P < 0.001.

### Ortholog and pathway sharing across species

We next examined orthogroup sharing across species (Fig. S5). Winter-elevated orthogroups showed greater conservation than winter-reduced: 27% of 330 winter-elevated orthogroups were shared across at least two species versus 21% of 470 winter-reduced orthogroups.

Variation in identified proteins partly reflects reference proteome completeness—*S. nutans*, with a draft genome containing 26% ambiguous bases, yielded fewer identifications (Table S2).

GO enrichment analysis revealed distinct functional signatures (Fig. S6a). Winter-elevated proteins were enriched for protein folding, stress response, cold acclimation, and ROS scavenging (Table S5). Winter-reduced proteins were enriched for microtubule-based processes, mitotic cell cycle, cytoskeleton organization, and cell wall biogenesis—processes associated with active growth (Table S6). The greater conservation of winter-elevated orthogroups suggests that protective responses to freezing are more constrained than growth-related processes active during summer. To identify the most conserved functional themes, we examined the 50 most winter-elevated proteins per species (Fig. 3; Table S7). LEAs and heat shock proteins (HSPs) were present in at least four of five species. A chi-square test on standard GO categories revealed significant species heterogeneity (χ²[70] = 97.5, *P* = 0.017), but GO terms may conflate cold-specific processes with broader stress responses (Barnes et al., 2023). 63% of proteins mapped to one of nine functional groups curated for relevance to cold (Fig. S6b). Heterogeneity remained significant (χ²[40] = 58.1, *P* = 0.032; Fig. 3a), indicating that while species share the same core functional categories, they differ in relative investment. Detoxification and ROS scavenging proteins were represented in all five species. Metabolism/osmoregulation proteins were less prominent in *A. gerardi*, while protein aggregation and membrane stability proteins were enriched in *A. gerardi* and *P. virgatum*. These patterns indicate a shared functional core with lineage-specific variation in relative allocation (Fig. 3a), particularly in metabolism/osmoregulation and protein folding/stability.

**Fig. 3.**
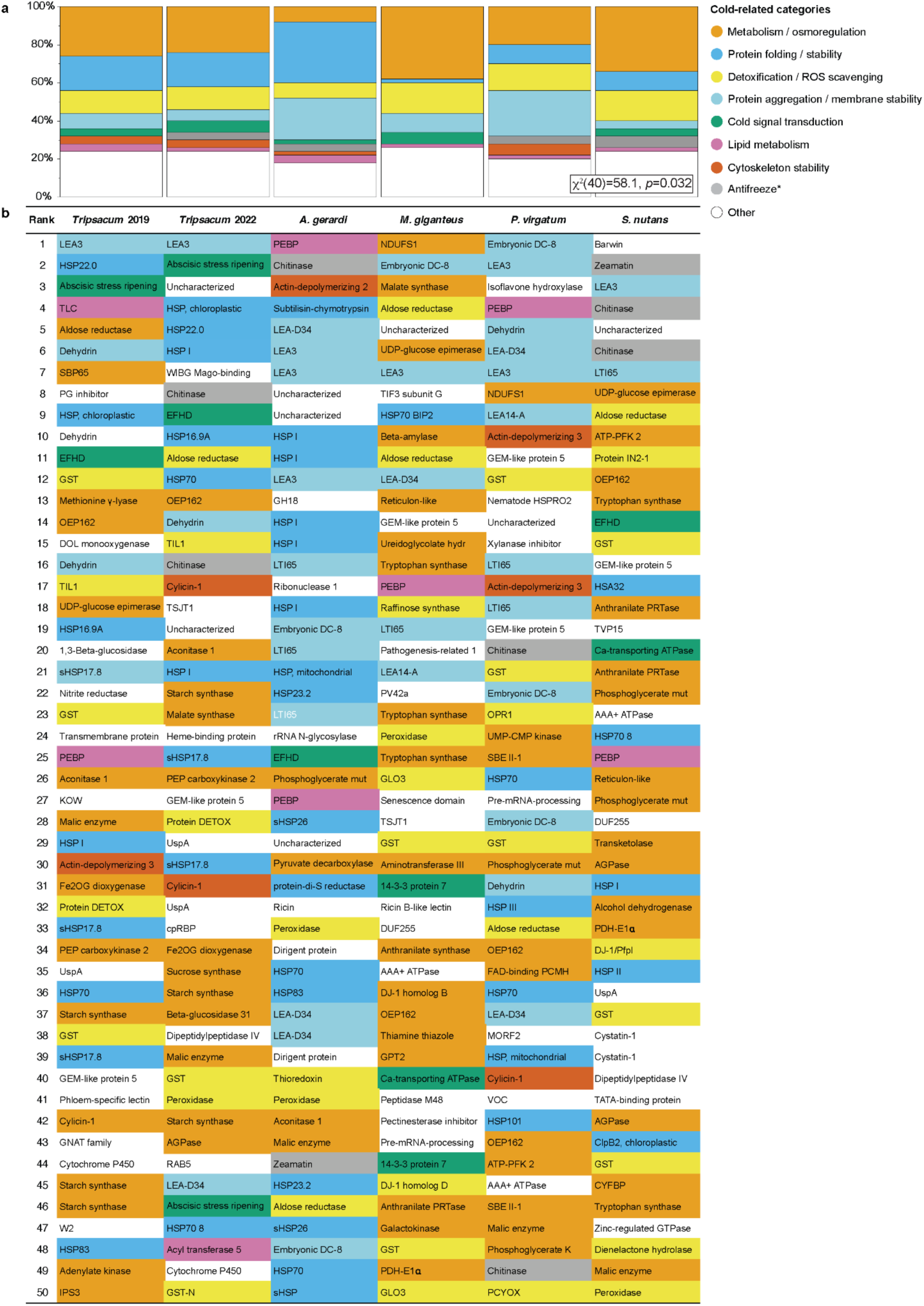
Overview of cold-related functional categories represented among the top 50 most winter-enriched rhizome proteins *Tripsacum dactyloides* and *T. floridanum* hybrids, *Andropogon gerardi*, *Miscanthus* × *giganteus*, *Panicum virgatum*, and *Sorghastrum nutans*. (a) Mosaic plot of cold-related functional categories across species. This plot visualizes the distribution of proteins among cold-related functional categories for each species. The relative abundance of each category within a species is shown, highlighting distinct functional profiles and species-specific variations in cold tolerance mechanisms. A Chi-square test for independence (χ²[40] = 58.1, *P* = 0.032) was performed to evaluate variation across species. **(b)** Visual representation of cold-related functional categories in rhizome proteins of five *Andropogoneae* species. This figure illustrates the top 50 most abundant proteins within the rhizomes of five *Andropogoneae* species. Proteins are classified into functional categories relevant to cold-related mechanisms, such as metabolism/osmoregulation, cold signal transduction, membrane stability, lipid metabolism, and others. Color-coding facilitates a comparative visualization of cold tolerance mechanisms across species. Antifreeze proteins are marked by an asterisk, as their classification is based solely on protein homology.

LEA3 (OG0022470) was the only ortholog ranked among the top 10 winter-elevated proteins in all five species and was the single most elevated protein in *Tripsacum* (log₂FC = 5.0; Fig. 3b; Table S7). Other functional categories showed species-specific representation: HSPs appeared in the top 10 for *Tripsacum* (HSP22, Class IV) and *S. nutans* (HSP23.2) but involved different family members; ROS scavenging was universal but employed glutathione transferases in four species and peroxidases in *A. gerardi*; putative antifreeze proteins (chitinases) ranked highly in *S. nutans* and *A. gerardi*. This pattern—a single conserved ortholog (LEA3) alongside functionally equivalent but distinct proteins in other categories—exemplifies parallel recruitment of LEA3 alongside functional convergence for chaperone and antioxidant functions.

### Evolutionary patterns in accumulated protein families

Phylogenetic analysis revealed distinct evolutionary patterns between LEAs and HSP families (Fig. 4). The control elongation factor 1 protein (EF1) family, previously shown to have stable expression at transcript and protein levels across conditions (Takamori et al., 2017, Fig. 4a), showed minimal seasonal accumulation and no phylogenetic clustering of abundance level patterns, consistent with their housekeeping function (Table S8). HSPs showed greater diversity (212 proteins across 37 orthogroups, Table S9) compared to LEAs (66 proteins in 20 orthogroups, Table S10), with control EF1 comprising 33 proteins in 4 orthogroups. Both LEA and HSP phylogeny (Figs. 4b and c) resolved into clear functional sub-families, but among HSPs the most strongly winter-elevated members (log₂FC > 2.5) belonged to different clades across different species, with only 32% of HSP proteins significantly elevated in winter. In contrast, LEA proteins (Fig. 4b) demonstrated more consistent winter abundance (47% of the proteins), particularly within the LEA3 subfamily, which displayed elevated abundance across all species despite having diverged earlier than other LEA clades.

**Fig. 4.**
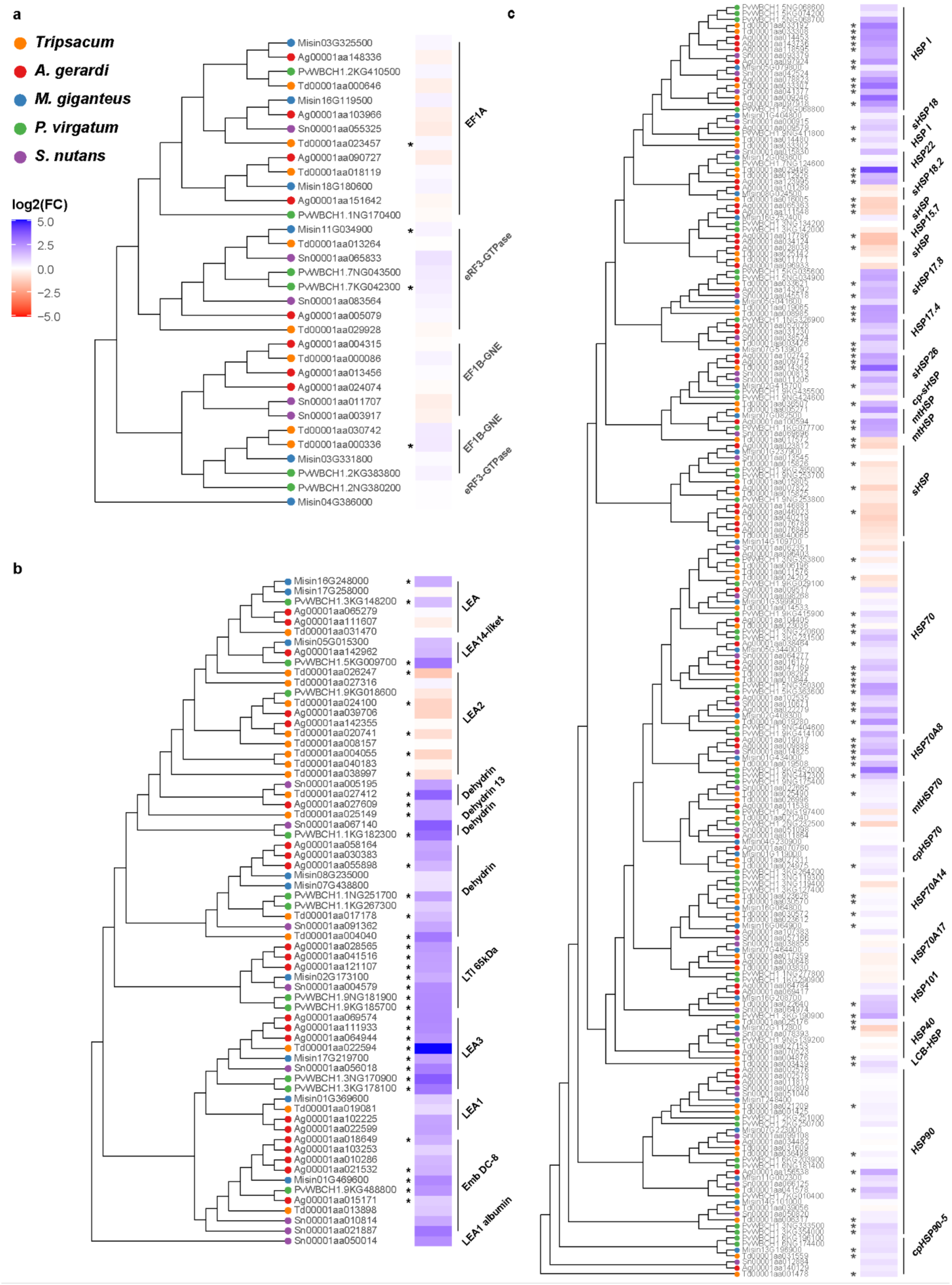
Phylogenetic analysis and seasonal abundance dynamics of selected protein families across *Tripsacum dactyloides* and *T. floridanum* hybrids, *Panicum virgatum*, *Andropogon gerardi*, *Miscanthus* × *giganteus*, and *Sorghastrum nutans*. Protein identifiers represent species-specific sequences, and brackets denote functional group annotations, merging adjacent orthogroups with similar functions for clarity. Heat maps illustrate log₂ fold-change (log₂FC) in protein abundance between winter and summer conditions, where blue indicates higher abundance in winter, and red signifies higher summer abundance. Panels show distinct protein families: **(a)** translation elongation factor 1 proteins (EF1; control group), **(b)** late embryogenesis abundant (LEA) proteins, and **(c)** heat shock proteins (HSPs). Asterisks (*) mark proteins exhibiting statistically significant differential accumulation (*P* < 0.05).

### Beyond rhizomes: Comparison with maize & Tripsacum mRNA

We next tested whether rhizome protein signatures corresponded to mRNA responses in other tissues. Cold-induced mRNA in F₂ seedling leaves of *Tripsacum*, derived from the same hybrids used in our rhizome study, showed weak but significant correlation with rhizome protein fold-changes (*ρ* = 0.29, *P* < 0.001; Fig. S7a). Other *Tripsacum* tissues showed weaker correlations (*ρ* = 0.08–0.16; Fig. S7b–e), as did maize orthologs (*ρ* = 0.01–0.09; Figs. S7f–i). Despite these weak overall correlations, LEA3 displayed strong cold induction in leaves of both species (log₂FC = 8 in *Tripsacum*, and 7 in maize; Fig. S7a and f). That maize induces LEA3 transcription to levels comparable to cold-tolerant *Tripsacum*, yet remains freezing-sensitive, suggests that transcriptional induction of *LEA3* alone does not confer protection.

### Structural and Functional Insights into LEA3: A Key Player in Freezing Tolerance

LEA3 (OG0022470) was the only ortholog consistently elevated across all five species, ranking as the top DAP in *Tripsacum* rhizomes (log₂FC = 5.0) and the most cold-induced transcript in leaves (log₂FC = 8). Sequence alignment revealed that all six PACMAD orthologs share the Group 3 LEA architecture (Fig. S8), featuring tandem 11-mer repeats that fold into amphipathic α-helices (Fig. 5b). LEA3 proteins are classified as intrinsically disordered proteins thought to adopt amphipathic alpha-helical conformations upon binding to membranes or other proteins during cellular dehydration (Kovacs et al. 2008). These repeats generate characteristic oscillations in Kyte-Doolittle hydropathy profiles (Fig. 5c; Table S11; Fig. S9), consistent with a hydrophobic face (positions 1, 2, 5, 9) and a charged hydrophilic face essential for desiccation and freezing tolerance (Dure, 1993). While the number of repeats varied—ranging from three (*M. giganteus*, *Z. mays*) to five (*T. dactyloides*, *A. gerardi*)—total LEA3 dosage was further modulated by gene copy number, with *A. gerardi* and *P. virgatum* accumulating multiple paralogs (Fig. 4b; Table S12). Notably, although *M. giganteus* and *Z. mays* share the same three-repeat architecture, their biophysical properties diverge. The *M. giganteus* ortholog retains hydrophilic consensus repeats, while the maize ortholog contains a variant third repeat (`AADAMEAAKQK`) with alanine and methionine substitutions that increase local hydrophobicity (repeat GRAVY −0.67 vs consensus mean −1.41; Table S12). This variation produces a distinct hydrophobic peak in the hydropathy profile (Fig. 5c) and a more pronounced hydrophobic face in the predicted structure (Fig. 5b), potentially compromising the balance between hydrophobic and hydrophilic faces required for effective cryoprotection. Hydrophobicity scores for individual LEA3 orthologs are provided in Table S11.

**Fig. 5.**
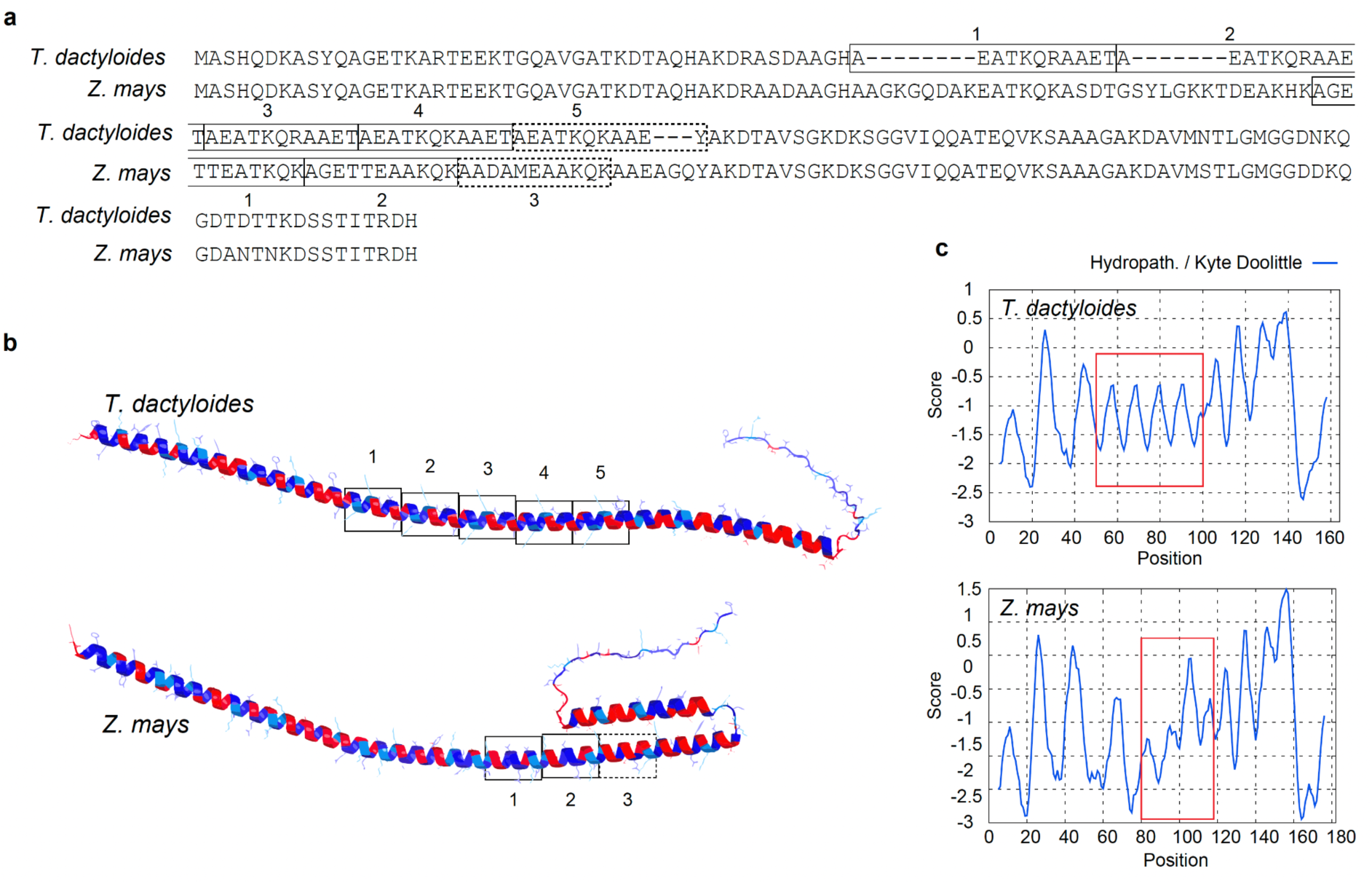
Protein sequence alignment, structure and hydrophobicity of LEA3 orthologs from *Tripsacum dactyloides* and *Zea mays*. **(a)** The alignment shows the LEA3 proteins from *T. dactyloides* (Td00001aa022594_T002) and, *Z. mays* (Zm00001eb294480), with tandem 11-mer repeat units highlighted and labeled numerically. *T. dactyloides* contains five repeat units (1–5); *Z. mays* contains three (1–3). The dashed box indicates a repeat unit with variant residues relative to the consensus repeat sequence. (**b**) Predicted secondary structure of LEA3 proteins using AlphaFold3, showing the conserved alpha-helical domains across *T. dactyloides* (top) and *Z. mays* (bottom). Repeat regions corresponding to labeled sequence motifs are boxed. Red and blue indicate regions of high hydrophilicity and hydrophobicity, respectively. **(c)** Kyte-Doolittle hydropathy plots for LEA3 proteins. The X-axis represents residue position, while the Y-axis indicates hydropathicity scores (positive values for hydrophobic regions, negative for hydrophilic regions). Red boxes highlight the amphipathic region encompassing the 11-mer tandem repeat array, corresponding to key domains in (b).

Hydropathy profiles from all cold-tolerant species examined show oscillating patterns consistent with amphipathic helix periodicity (Fig. S9). Species with more repeats exhibit more oscillation cycles and consequently longer amphipathic regions. Outgroup LEA3 proteins from wheat (TdLEA3; Koubaa et al., 2019) and *Arabidopsis* (AtLEA7; Popova et al., 2015) showed comparable repeat architecture. These observations suggest that transcriptional induction of *LEA3* alone may not ensure freezing tolerance and that functional protection may depend on both protein accumulation in the appropriate tissue and the biophysical properties of the tandem repeats of the 11-mer motif.

## Discussion

Our analysis of rhizome proteomes from five independently cold-adapted PACMAD grasses reveals that protein-level cold responses combine ancestral retention of core protective orthologs with convergent deployment of functionally equivalent but phylogenetically distinct proteins. The clearest example of retention is LEA3, accumulated across all five lineages: its early divergence from other LEA subfamilies (Fig. 4b) and the conservation of its tandem 11-mer repeat architecture across cold-tolerant species suggest that the cryoprotective capacity of LEA3 predates the PACMAD radiation and has been maintained under constraint in lineages that colonized cold environments (Fig. 1e). In contrast, the chaperone response illustrates convergent evolution: different HSP family members are recruited in different species, suggesting that chaperone-mediated protection permits greater flexibility in which family members are deployed. To our knowledge, this study provides the first quantitative comparison of cross-species fold-change conservation at the protein level in a freezing-stress context, complementing transcript-level comparisons in *Pooideae* (Schubert et al., 2019) and desiccation-tolerance studies in resurrection grasses (Marks et al., 2024).

The most striking finding is the evolutionary constraint on response magnitude. Among shared cold-responsive orthogroups, cross-species fold-change correlations were substantially higher (*ρ* = 0.80, Fig. 2b) than among background proteins (*ρ* = 0.45; *P* = 1.1 × 10⁻⁵), indicating that when freezing tolerance evolved independently across PACMAD lineages, it proceeded through quantitatively similar responses in a core set of protective proteins. Our common-garden design ensured identical winter conditions across all genotypes, with winter collections following sustained sub-zero exposure reaching −16°C and soil temperatures below 2–5°C for 20–22 days (Brotzge et al., 2020), while summer collections captured vigorous growth with actively emerging rhizome buds after >90 mm cumulative precipitation (Fig. S1), establishing these seasonal contrasts as cold-driven rather than drought-related. A limitation of this design is that all species experienced identical winter conditions rather than stress calibrated to each species’ tolerance threshold.

With these responses established as cold-driven, their conservation contrasts sharply with transcript-level patterns: comparative transcriptomics in *Pooideae* found generally low similarity in cold-responsive gene expression across species (*r* = 0.09–0.40; Schubert et al., 2019), with nearly half of all cold-responsive genes being species-specific. This comparison spans different clades, tissues, and experimental designs — our PACMAD rhizome proteomes versus *Pooideae* seedling transcriptomes — so the gap between protein-level and transcript-level correlations may be confounded by these differences. Nevertheless, the direction is consistent with the framework of Cope et al. (2025), who demonstrated that protein abundances are under stronger stabilizing selection than mRNA abundances over macroevolutionary timescales, with mutations affecting mRNA levels largely buffered before they alter protein output. If this buffering operates on cold-responsive genes as it does genome-wide, then transcript-level comparisons would systematically underestimate the degree of evolutionary conservation in stress responses. Notably, this constraint applies specifically to the dynamic response: absolute abundance correlations for shared cold-responsive proteins were lower than background in both summer (*ρ* = 0.62 vs 0.68; *P* = 0.004) and winter (Fig. 2b), indicating that these proteins are more divergent across species in their resting and stress-state levels. What is conserved is not the absolute abundance in either season, but the magnitude of accumulation when freezing arrives. This divergence in absolute levels likely reflects the distinct physiological contexts of each species’ rhizomes, including differences in dormancy depth, metabolic rate, or tissue composition, while the conserved fold-change suggests that selection acts primarily on the dynamic response rather than on any single optimal protein concentration.

LEA3 provides the clearest illustration of this constraint and connects the evolutionary pattern to a mechanistic hypothesis. LEA proteins function as molecular shields during freezing through membrane stabilization, the prevention of protein aggregation, and the protection of client proteins through steric exclusion and stress induced conformational changes (Kovacs et al., 2008; Cuevas-Velazquez et al., 2017). These functions depend on the repeating 11-mer motif architecture, described by Dure (1993), in which apolar residues form the hydrophobic face of α-helices that bind membranes and partially unfolded proteins during dehydration and freezing (Koubaa *et al*., 2019; Singh & Graether, 2021).

The structural data support a model in which cryoprotective efficacy arises from three factors: amphipathic helix length per protein (determined by repeat number), total protein abundance (increased through paralog accumulation in *A. gerardi* and *P. virgatum*), and repeat composition. All cold-tolerant species and outgroups maintain conserved amphipathic character across their 11-mer repeats (Figs. S8, S9; Table S12), whereas the variant third repeat of Z. mays shows an exaggerated hydrophobic face that may compromise the amphipathic balance required for effective cryoprotection (Table S12). AlphaFold3 multimer predictions reinforce this distinction: while both Td and ZmLEA3 form tetrameric assemblies, they adopt distinct packing geometries (Fig. S10), consistent with observations that cereal LEA3 proteins require conserved domains for oligomerization (NDong *et al*., 2002) and Arabidopsis LEA3 proteins form higher-order assemblies under stress (Ginsawaeng et al., 2021; Romero-Pérez et al., 2024). The variant hydrophobic face in ZmLEA3 may disrupt the optimal packing arrangement required for effective cryoprotection. This structural divergence helps explain why, although ZmLEA3 is cold-inducible and confers partial cold tolerance (Liu et al., 2013), maize remains freezing-sensitive despite accumulating transcripts to levels comparable to the cold-tolerant *Tripsacum* in leaves (Fig. S7f). Transcriptional induction alone does not ensure protection; functional efficacy depends on both sufficient abundance and activity, the latter shaped by the biophysical properties of the repeat region. The early divergence of LEA3 from other LEA subfamilies (Fig. 4b) suggests that the molecular properties enabling cryoprotection were likely present before the PACMAD radiation. Cold-tolerant lineages maintained or expanded LEA3 repeat number and in some cases accumulated multiple paralogs, while in maize — where freezing selection was relaxed — repeat number was not expanded and the third repeat diverged from the hydrophilic consensus.

Yet, freezing survival involves the coordinated action of numerous protective mechanisms, and restoring a single protein’s architecture in a freezing-sensitive species would be unlikely to confer full tolerance. Rather, LEA3 illustrates how structural constraints at the protein level could limit the efficacy of otherwise conserved transcriptional responses.

More broadly, several of the most consistently accumulated proteins across PACMAD rhizomes have well-characterized orthologs implicated in cold acclimation in *Arabidopsis*, including LEA proteins, small HSPs, and glutathione S-transferases (Marrs, 1996; Sun et al., 2002; Hundertmark & Hincha, 2008), indicating deep conservation of core protective pathways across monocots and dicots. However, some highly accumulated proteins in PACMAD rhizomes—notably PEBP family members and certain chitinase-like proteins—have not been prominently featured in *Arabidopsis* cold studies, potentially reflecting grass-specific or rhizome-specific adaptations (Roeder et al., 2025).

The presence of ABA-inducible proteins and dehydrins among the most accumulated winter proteins across species is consistent with shared physiological pathways between drought and frost responses. While evidence shows that drought tolerance evolved before frost tolerance in PACMAD grasses, suggesting they may have been co-opted, the extent to which this actually happened remains unclear (Schat et al., 2025). In contrast, Pooideae grasses show a different trajectory, where frost tolerance appears to have evolved independently of, or even prior to, drought tolerance (Stolsmo et al., 2024; Schat et al., 2025).

In conclusion, rhizome freezing tolerance across independently cold-adapted PACMAD grasses is governed by the retention of ancestral protective proteins under quantitative constraint on response magnitude, combined with convergent deployment of diverse protein families for chaperone and antioxidant functions. This constraint is substantially stronger at the protein level than previously observed at the transcript level. The differences in LEA3 repeat number and composition between cold-tolerant PACMAD species and maize offer a candidate mechanism for enhancing freezing resilience in maize. Future work should prioritize functional validation of LEA3 repeat variants, particularly testing whether restoring the ancestral hydrophilic consensus or introducing additional paralogs can enhance frost tolerance in maize.

## Supporting information

Supplemental Codes

## Acknowledgments

We thank Cornell Botanic Gardens for their support of this study and for maintaining the field plots that provided essential plant material, particularly Kristine Boys, Emily Detrick (Elizabeth Weaver Director of Horticulture), and Todd Bittner (Director of Natural Areas). We acknowledge the Proteomics and Metabolomics Facility at the Cornell Institute of Biotechnology (RRID:SCR_021743) for providing mass spectrometry services and data analysis. We are grateful to Prof. Hening Lin for funding support from the HHMI Transformative Technology 2019 program for the Orbitrap Eclipse system, and to Dr. Sheng Zhang for NIH SIG grant support (1S10 OD017992-01 for the Orbitrap Fusion mass spectrometer and 1S10 RR025449-01 for the Orbitrap Velos/Elite mass spectrometer). We extend our appreciation to Elizabeth T. Anderson and Robert W. Sherwood at the Proteomics and Metabolomics Facility for their technical expertise and assistance with sample processing and analysis. We thank the Genomics Facility (RRID:SCR_021727) of the Biotechnology Resource Center at the Cornell Institute of Biotechnology for their assistance with sequencing experiments. Additional computational resources and data management were provided by the Bioinformatics Facility (RRID:SCR_021757) at the Cornell Institute of Biotechnology. This research is made possible by the New York State (NYS) Mesonet. Original funding for the NYS Mesonet (NYSM) buildup was provided by the Federal Emergency Management Agency grant FEMA-4085-DR-NY. The continued operation and maintenance of the NYSM is supported by the National Mesonet Program, University at Albany, Federal and private grants, and others. We acknowledge all who have contributed to the preservation, cultivation and study of wild and domesticated grasses.

## Author Contributions

M.C.R., T.E.R., and E.O. conceived and designed the study. N.L. and T.E.R. developed the long-term sampling strategy that enabled seasonal comparisons. N.L., T.E.R., T.L., S.-K.H., C.M.G., and E.O. performed sample collection and experimental work. T.L., R.A., and Q.F. conducted proteomics sample preparation, LC-MS/MS, and data processing. C.M.G., J.H., T.L., and J.Z. performed RNA-seq analysis. J.Z., S.-K.H., and E.O. carried out bioinformatics and statistical analysis. J.Z., S.-K.H., C.O.H., and E.O. prepared the figures. E.O., J.Z., L.J.B., and E.S.B. wrote the manuscript. C.O.H., L.J.B., S.-K.H., J.H., Q.F., E.S.B., and M.C.R. reviewed and edited the manuscript. E.O. and M.C.R. supervised the project and E.S.B. and M.C.R. acquired funding.

## Data Availability

The mass spectrometry proteomics data have been deposited to the ProteomeXchange Consortium via the PRIDE (Perez-Riverol et al., 2025) partner repository with the dataset identifier PXD063668 (Sample key file in Table S13). RNA-seq data are available under BioProject accession numbers PRJNA705456 (*Zea mays*) and PRJNA1260937 (*Tripsacum dactyloides*) in the NCBI BioProject database (https://www.ncbi.nlm.nih.gov/bioproject/). Genome assemblies and annotations used in this study are publicly available from MaizeGDB (Woodhouse et al., 2021) at https://maizegdb.org/PanAnd_project for *Tripsacum dactyloides* (Td-FL_9056069_6-DRAFT-PanAnd-1.0), *Andropogon gerardi* (Ag-CAM1351-DRAFT-PanAnd-1.0), and *Sorghastrum nutans* (Sn-CAM1369-DRAFT-PanAnd-1.0), and from the JGI Data Portal (Goodstein et al., 2012) at https://data.jgi.doe.gov/ for *Miscanthus* × *giganteus* (Msinensis_497_v7.0) and *Panicum virgatum* (Pvirgatumvar_WBCHAP1_778_v1.0), along with their corresponding GFF annotation files. Functional annotations, data tables, and analysis scripts are available from GitHub at https://github.com/elor77/PACMAD-Rhizome-Proteomics.

## Supporting Information

The following Supporting Information is available for this article:

**Fig. S1.** Rhizome morphology at sampling.

**Fig. S2.** Principal component analysis of log-transformed protein abundance data.

**Fig. S3.** Biological processes (BPs) within curated cold-related functional categories.

**Fig. S4.** Volcano plots showing differentially accumulated proteins in sampled species.

**Fig. S5.** Upset plots of shared differentially accumulated proteins among species in winter and summer.

**Fig. S6.** Enriched BPs in rhizomes of five PACMAD grass species and BP distribution within curated cold-related functional categories.

**Fig. S7.** Scatterplots showing the relationship between rhizome proteins and cold-responsive mRNA expression in *Tripsacum* and maize tissues.

**Fig. S8.** MAFFT alignment of LEA3 across six PACMAD species.

**Fig. S9.** Hydropathy profiles of LEA3 orthologs.

**Fig. S10.** AlphaFold 3 prediction of LEA3 homo-tetramer assemblies in *Tripsacum* and *Zea mays*.

**Table S1.** Normalized protein abundances (log_2_-transformed) for all samples.

**Table S2.** Genome assembly statistics for the five species analyzed.

**Table S3.** List of 4,888 orthogroups identified across species.

**Table S4.** Log_2_ fold-changes (Winter/Summer) for all quantified proteins.

**Table S5.** GO enrichment analysis for winter-upregulated proteins.

**Table S6.** GO enrichment analysis for winter-downregulated proteins.

**Table S7.** Top 50 most abundant proteins in winter vs summer.

**Table S8.** Elongation Factor 1-alpha (EF1) orthologs and abundance data.

**Table S9.** Heat Shock Protein (HSP) orthologs and abundance data.

**Table S10.** LEA3 orthologs and abundance data.

**Table S11.** Hydrophobicity scores for LEA3 orthologs.

**Table S12.** Biophysical properties of individual 11mer LEA3 (OG0022470) tandem repeats.

**Table S13.** PRIDE sample key and metadata.

**Methods S1** Detailed protocols for protein extraction, S-Trap digestion, LC–MS/MS search parameters, RNA-seq alignment settings, and bioinformatics pipeline parameters.

**Code S1** R script for importing original Ag/Mg multiplexed peptide intensities.

**Code S2** R script for re-aligning peptides using DIAMOND.

**Code S3** R script for summarizing peptides into protein-level data.

