## Supplemental Methods S1 for "Constrained evolution of a core winter proteome across independently cold-adapted PACMAD grasses"

**Supporting Information Methods S1**

Detailed environmental data, protocols for protein extraction, S-Trap digestion, LC–MS/MS search parameters, proteins abundance statistical analyses, GO enrichment, RNA-seq alignment settings, and bioinformatics pipeline parameters for: Constrained evolution of a core winter proteome across independently cold-adapted PACMAD grasses by Oren et al.

*Sampling environmental data*

In 2019, summer samples were collected on August 11 following 121 mm of precipitation in the preceding 30 days, and mean soil temperature at 5 cm depth was 19.9°C. Winter samples were collected on January 8, 2020, after 22 days with soil temperature below 2°C. In 2022, summer sampling occurred on August 15 following 93 mm of precipitation; 30-day mean soil temperature was 20.6°C. Winter sampling on December 9 followed 20 days with soil temperature below 5°C.

*Protein Extraction*

Fresh rhizome tissue was ground in liquid nitrogen. The ground sample was suspended in a lysis buffer composed of 1× PBS (pH 8), 10 mM EDTA, 900 mM sucrose, and 0.4% β-mercaptoethanol. The suspension was vortexed and incubated on ice, then mixed with an equal volume of phenol buffered to pH 8 with Tris-EDTA (TE). After centrifugation and phase separation, the phenol phase containing proteins was collected. Proteins were precipitated from the phenol phase using ice-cold ammonium acetate/methanol solution (100 mM w/v ammonium acetate in 100% methanol) with protease and phosphatase inhibitors. After precipitation the pellets were washed five times with three different solutions and frozen.

*S-Trap Digestion*

Protein pellets were dissolved in 8 M urea buffer. Protein concentration was determined using a BCA protein assay. A set amount of each protein sample underwent S-Trap digestion, including treatment with dithiothreitol (DTT) for reduction and iodoacetamide (IAA) for alkylation. The sample underwent multiple rounds of centrifugation with the addition of 8 M urea, followed by 50 mM ammonium bicarbonate (ABC) or triethylammonium bicarbonate (TEAB). The sample was digested using a trypsin buffer, centrifuged, and then dried by vacuum freeze-drying for storage prior to subsequent TMT labeling.

*High-pH Reversed-Phase Fractionation*

Labeled peptides were fractionated by high-pH reversed-phase chromatography on a Dionex Ultimate 3000 HPLC system. A total of 48 fractions were collected and concatenated into 10 pools using a meandering pooling scheme designed to maximize orthogonality with subsequent low-pH nanoLC separation.

*Database Search Parameters*

Samples were injected into a nanoLC system coupled to an Orbitrap Eclipse mass spectrometer using an RTS-SPS-$MS^3$ method (Fu *et al.*, 2021). Raw MS data were processed in Proteome Discoverer 2.5 using the Sequest HT search engine against species-specific reference proteomes (see main text, Peptide Mapping and Reference Proteome Assignment). Trypsin was specified as the proteolytic enzyme, allowing up to two missed cleavages with a minimum peptide length of six amino acids. Fixed modifications included TMTpro tags on peptide N-termini and lysine residues and carbamidomethylation of cysteine. Dynamic modifications included methionine oxidation, asparagine/glutamine deamidation, N-terminal acetylation, methionine loss, and methionine loss with acetylation. Precursor mass tolerance was set to 10 ppm and fragment mass tolerance to 0.6 Da.

*Peptide and Protein Validation*

Peptide spectral matches were filtered using Percolator with a strict target FDR of 0.01 and a relaxed target FDR of 0.05. Protein identifications were validated using the Protein FDR Validator node with the same strict (0.01) and relaxed (0.05) FDR thresholds. Proteins were initially identified with a minimum of one unique peptide and subsequently filtered to retain only those supported by at least two distinct peptides for downstream analysis.

*Reporter Ion Quantification*

Protein quantification was based on the summed reporter ion intensities of unique and razor peptides, filtered with a minimum signal-to-noise ratio of 10, SPS mass matches ≥65%, and a co-isolation threshold of 50%. No minimum channel occupancy threshold was applied, allowing quantification of proteins detected in subsets of samples. Total peptide amount normalization was applied across TMT channels. Protein abundance ratios were calculated based on summed reporter ion intensities, and hypothesis testing was performed using one-way ANOVA within Proteome Discoverer 2.5.

*Protein abundance normalization and statistical analysis*

PCA was performed using the FactoMineR::PCA function in R with centering and scaling. For cross-species comparison at the orthogroup level, in cases where multiple proteins belonged to the same orthogroup within a species, $log_2$ fold-change values were averaged to generate a single orthogroup-level fold change. For differential abundance analysis, only proteins quantified in both winter and summer samples within each species were retained. All statistical analyses and visualization were performed in R version 4.4.3, using RStudio 2024.12.1+563 ("Kousa Dogwood" release).

*Functional annotation and GO enrichment*

Orthogroups served as the basis for cross-species comparisons and functional propagation of annotations, particularly for proteins with limited annotation coverage outside *T. dactyloides*. For non-significant proteins, functional annotations were propagated from *Tripsacum dactyloides* to orthologous proteins within the same orthogroup when available. Remaining unannotated entries were manually curated based on sequence similarity and conserved domain features.

The topGO R package (v2.50.0; Alexa & Rahnenfuhrer, 2017), “elim” algorithm with Fisher’s exact test was used to reduce redundancy and account for GO hierarchy. The background set included all proteins quantified across the experiment. The top 15 enriched Biological Process (BP) terms in each group were visualized using dot plots in ggplot2, incorporating log-transformed P-values, gene ratios, and counts of significant proteins.

*Phylogenetic Analysis of Candidate Protein Families*

Proteins for phylogenetic trait mapping were identified by text-mining functional annotations produced by PANNZER2 for both the *Tripsacum dactyloides* proteome and the DAP sets from all five species, searching specifically for GO terms associated with LEA proteins, HSPs, and EF1. Extracted orthogroup identifiers from our $\log_2$ fold-change matrix were then used to retrieve the corresponding protein IDs, which served as input for sequence extraction. Full-length amino-acid sequences were obtained from each species' reference proteome using seqkit grep (v2.0) with protein headers matching extracted IDs. Multiple sequence alignments were generated in MAFFT (v7.475; (Katoh & Standley, 2013) under default settings. Maximum-likelihood trees were inferred using raxmlHPC (RAxML-8.2.13; (Stamatakis, 2014) with the `PROTGAMMAJTT` substitution model and 100 rapid bootstraps. Each resulting tree was rooted on a biologically appropriate outgroup (e.g., a chloroplastic class I small HSP) and visualized in R (v4.1.0) using the ggtree package.

*RNA-seq Processing Parameters*

Paired-end RNA-seq reads were quality-filtered and trimmed using Trimmomatic (v0.32; (Bolger *et al.*, 2014) to remove Illumina adapter bases, discarding leading bases with a phred score below 35 and trailing bases with a phred score below 20. Reads shorter than 36 bp after trimming were discarded.

Trimmed reads were aligned to the Tripsacum dactyloides genome (Td-FL_9056069_6-DRAFT-PanAnd-1.0.fasta) and annotation (Td-FL_9056069_6-DRAFT-PanAnd-1.0_final.gff3) using STAR v2.7.10b in two-pass mode (`--twopassMode Basic`), allowing each read to map to a maximum of 20 genomic locations (`--outFilterMultimapNmax 20`). Gene-level counts were extracted from uniquely mapped reads using featureCounts v2.0.3 with default settings.

Differential gene expression was assessed using DESeq2 (Love *et al.*, 2014), modeling both genotype and temperature as main effects (~ genotype + temperature). Contrasts were made between 5°C and 22°C within each of eight $F_2$ families, with 5°C as the reference level. Raw counts were normalized using the estimateSizeFactors method. Genes were considered differentially expressed at adjusted $P < 0.05$ (Benjamini-Hochberg correction) and $|\log_2 FC| > 1$.
