## Supplemental Figures for "Constrained evolution of a core winter proteome across independently cold-adapted PACMAD grasses"

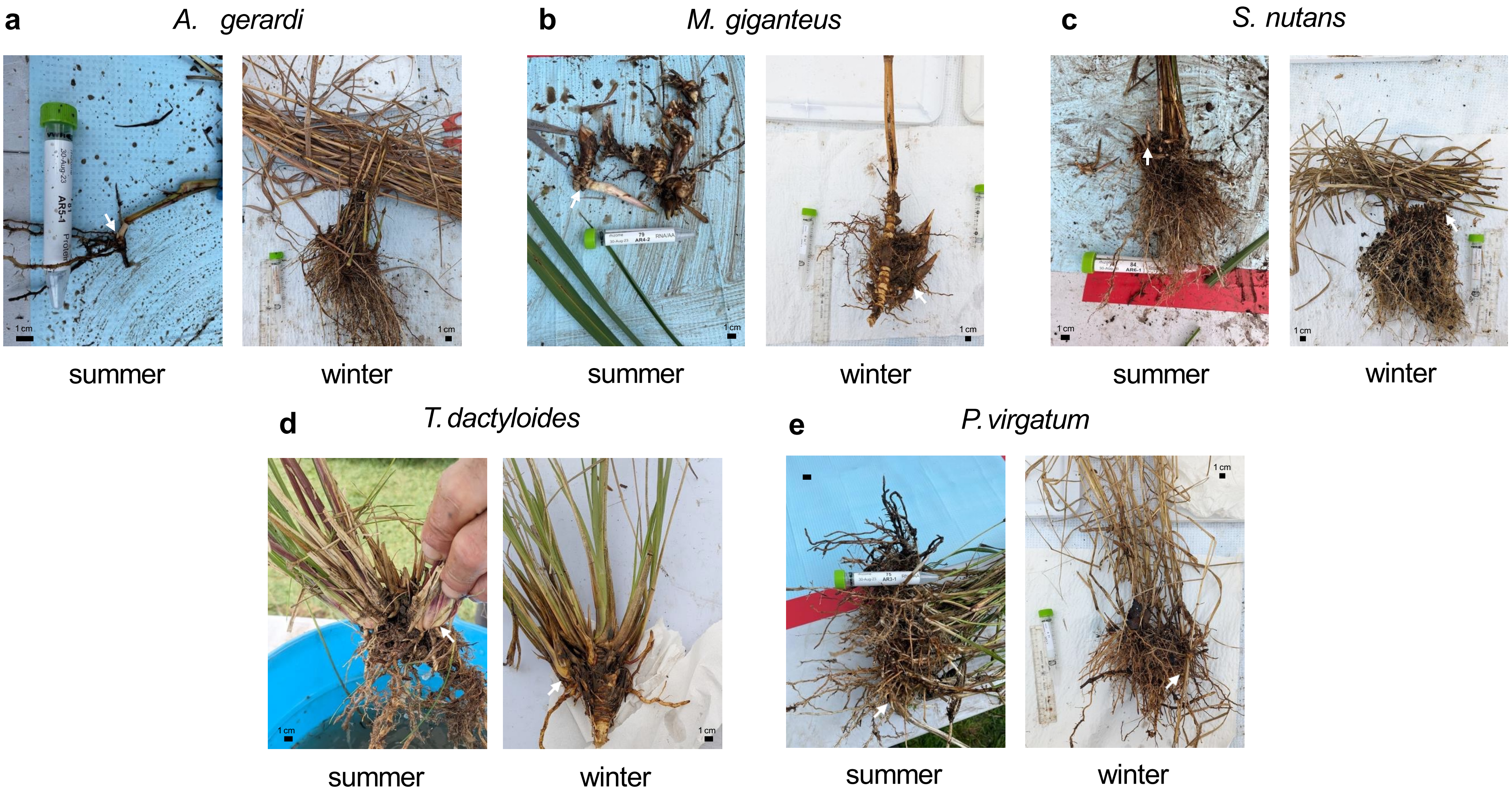


**Fig. S1. Rhizome morphology at sampling.** Paired images display representative rhizome tissues (white arrows) collected during summer (August, left) and winter (January, right) for **(a)** *Andropogon gerardi*, **(b)** *Miscanthus* × *giganteus*, **(c)** *Sorghastrum nutans*, **(d)** *Tripsacum dactyloides* and *T. floridanum* hybrids, and **(e)** *Panicum virgatum*. Summer samples exhibit active bud growth and pale storage tissue. Winter samples exhibit dormancy and epidermal hardening. Proteomic analysis was restricted to mid-section of these storage organs. Scale bars represent approx. 1 cm.

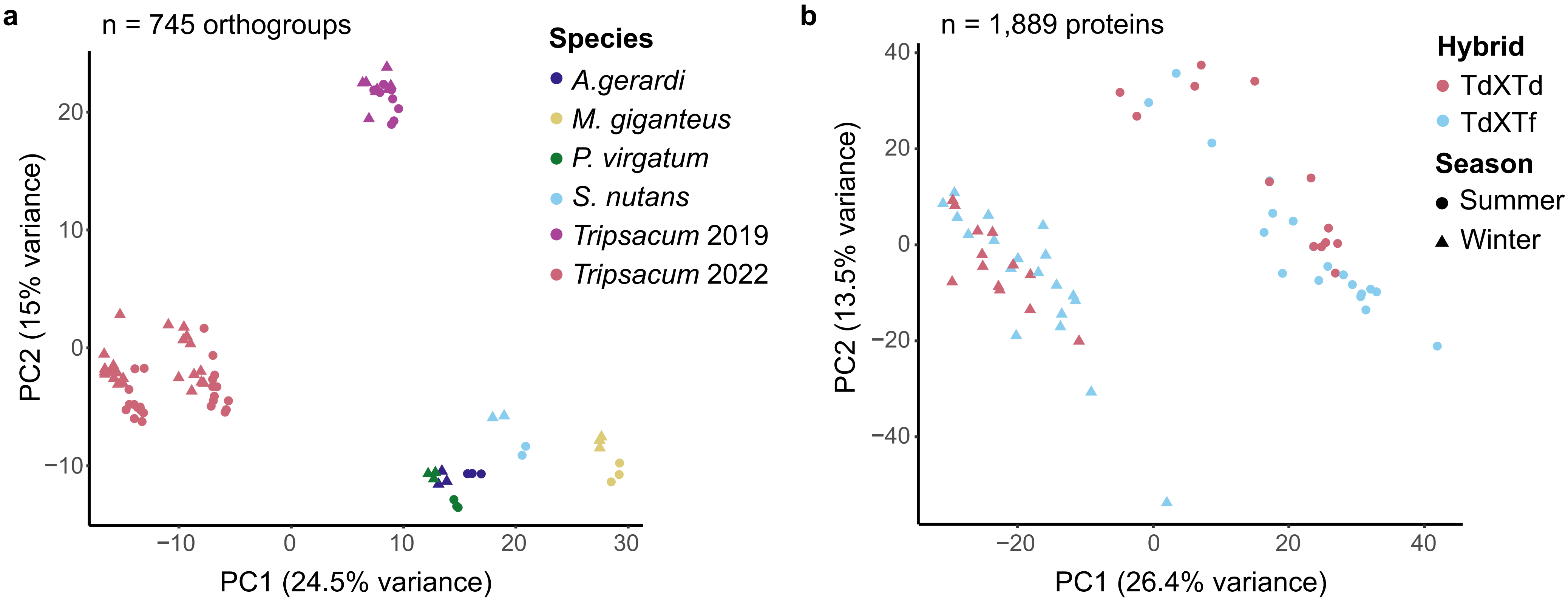


**Fig. S2. Principal component analysis (PCA) of log-transformed protein abundance data. (a)** PCA of 745 orthogroups from five species *Tripsacum dactyloides and T. floridanum hybrids* (Td-2019, Td-2022), *Andropogon gerardi* (Ag), *Miscanthus × giganteus* (Mg), *Panicum virgatum* (Pv), and *Sorghastrum nutans* (Sn). **(b)** PCA of 1,889 proteins from *Tripsacum* hybrids. Points are colored by hybrid type–intraspecific *T. dactyloides* hybrids (Td × Td) and interspecific *T. dactyloides × T. floridanum* hybrids (Td × Tf). Shapes represent season (circle = summer, triangle = winter). The percentage of total variance explained by each principal component is shown on the axes.

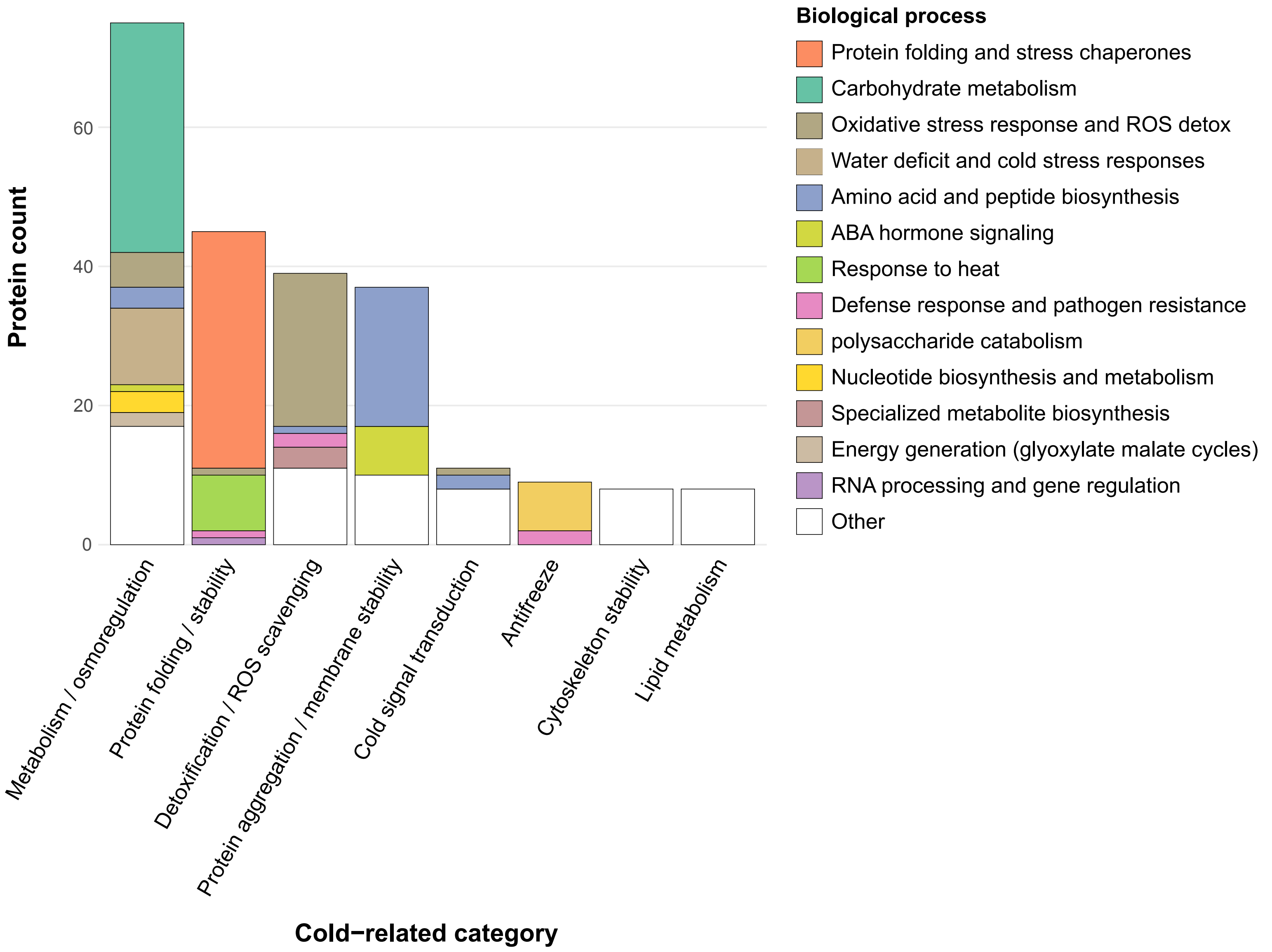


**Fig. S3 Stacked bar chart illustrating the composition of biological processes (BPs) within curated cold-related functional categories**. The plot includes 230 proteins from the top 50 differentially accumulated proteins (DAPs) accumulated in winter rhizomes of *Tripsacum dactyloides* and *T. floridanum* hybrids, *Andropogon gerardi*, *Miscanthus* × *giganteus*, *Panicum virgatum*, and *Sorghastrum nutans*, after excluding those assigned to the cold response category "Other" (i.e., proteins with annotations deemed less relevant to cold tolerance interpretation). Cold-related categories are shown on the x-axis and sorted by total protein count. Each bar is color-coded by BP, highlighting the functional composition of each category. Less informative or poorly annotated BPs were grouped under "Other" in the legend. This visualization supports the manual consolidation of specific biological processes into broader cold-response categories used throughout the study.

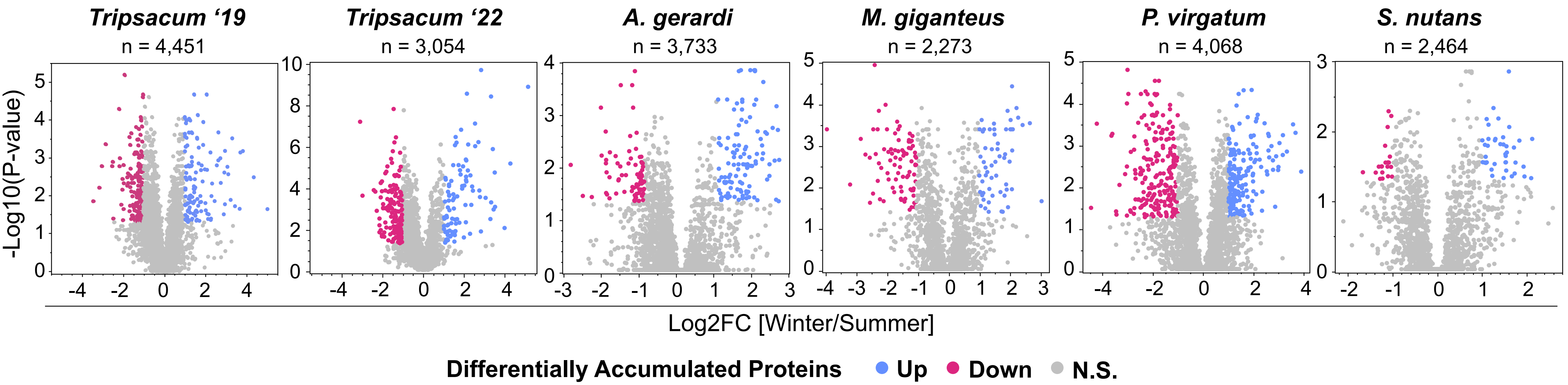


**Fig. S4.** Volcano plots showing differentially accumulated proteins (DAPs) in rhizomes of *Tripsacum dactyloides and T. floridanum hybrids, Andropogon gerardi, Miscanthus* × *giganteus*, *Panicum virgatum*, and *Sorghastrum nutans* under winter vs. summer conditions, with total number of proteins identified listed. Proteins with significantly elevated (blue) and reduced (pink) abundance in winter relative to summer are highlighted, with non-significant proteins in gray. The y-axis represents the significance as $-\log_{10}$(p-value), and the x-axis shows the $\log_2$ fold-change ($\log_2$FC) of the DAPs between winter and summer.

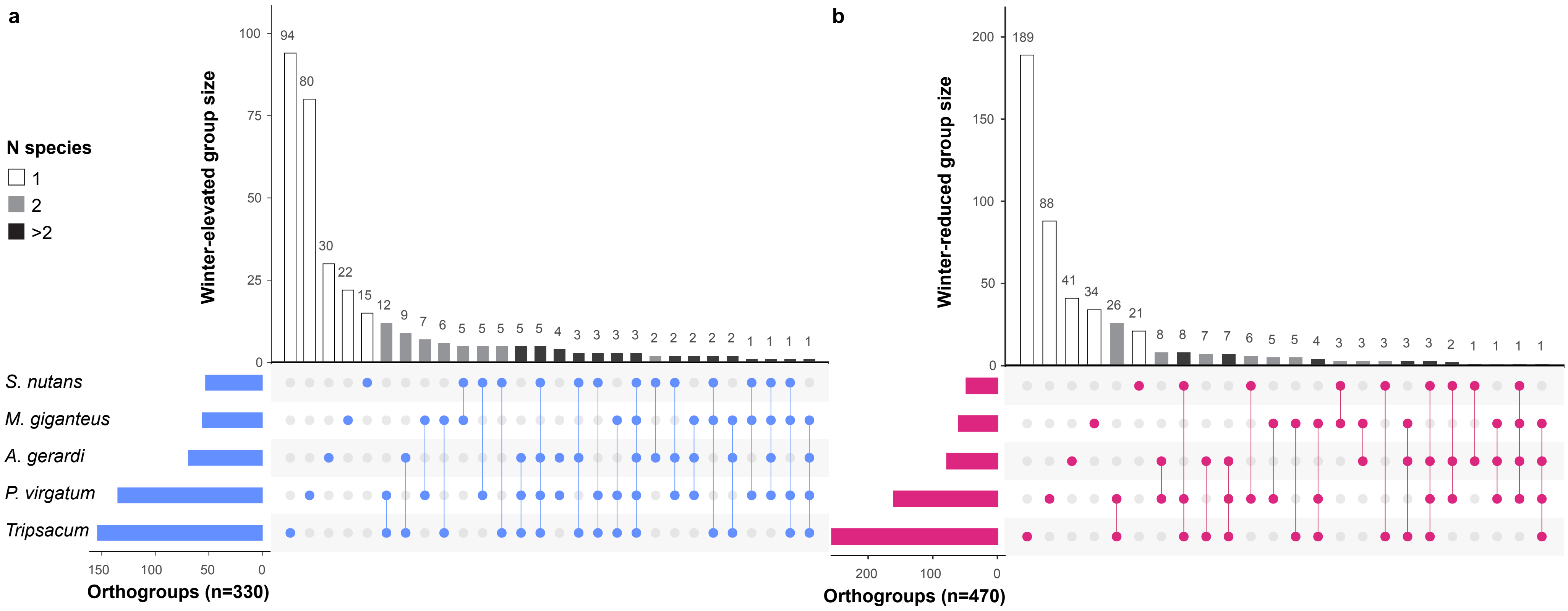


**Fig. S5. Orthogroup sharing of significantly DAPs among *Tripsacum dactyloides and T. floridanum hybrids, Andropogon gerardi, Miscanthus* × *giganteus*, *Panicum virgatum*, and *Sorghastrum nutans*.** **(a)** Winter-elevated. **(b)** Winter-reduced. Set sizes (horizontal bars) indicate the total number of differentially abundant orthogroups identified per species. Intersection sizes (vertical bars) represent orthogroups unique to individual species or shared among multiple species. Bar shading indicates the number of species sharing each orthogroup: unique to one species (white), shared between two species (gray), or shared among more than two species (black). Dots below the plot illustrate species-specific orthogroup intersections.

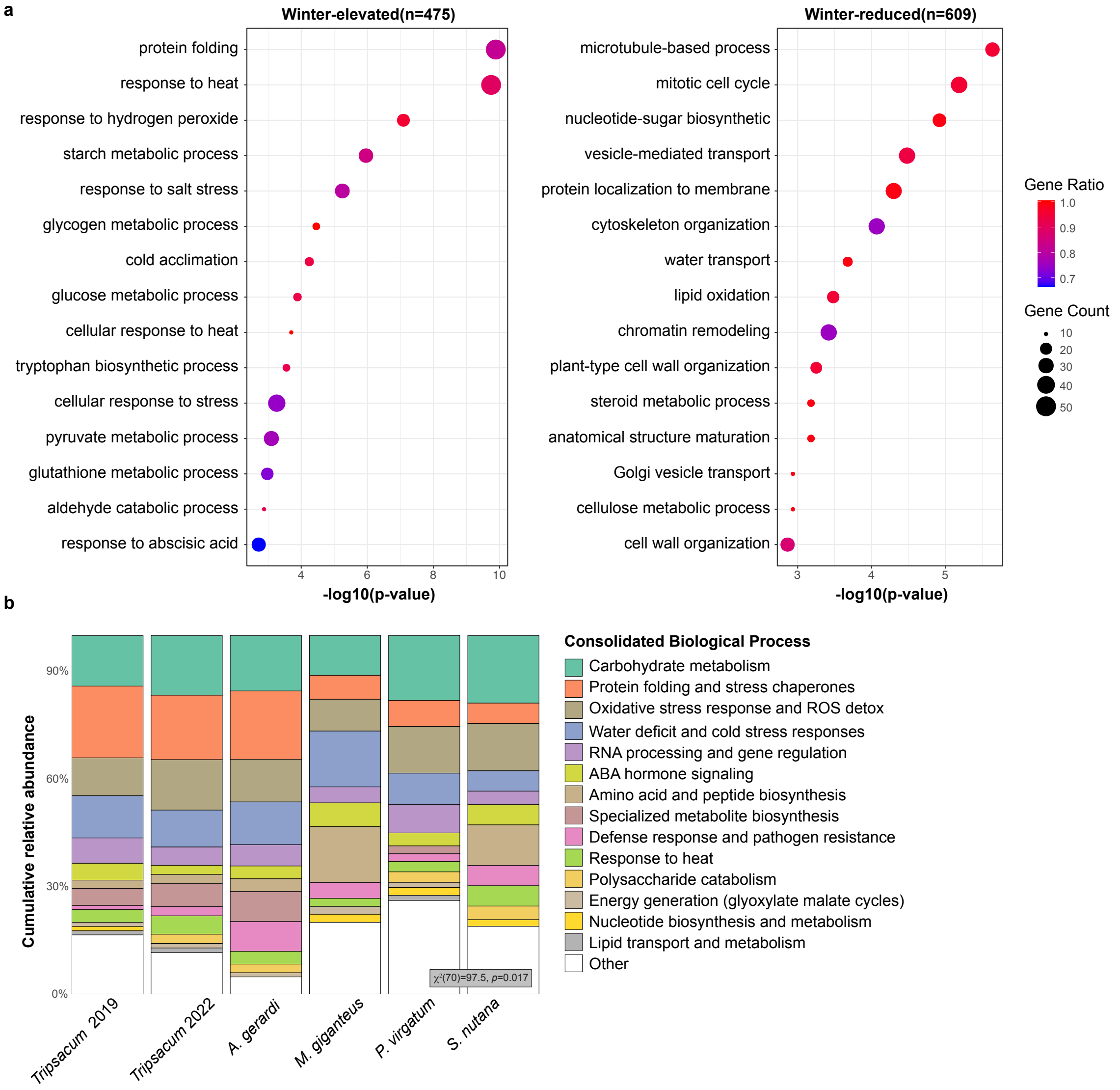


**Fig. S6 (a)** Enriched biological processes (BP) of *Tripsacum dactyloides* and *T. floridanum* hybrids, *Andropogon gerardi*, *Miscanthus × giganteus*, *Panicum virgatum*, and *Sorghastrum nutans* rhizomes among differentially accumulated proteins (DAPs) in 475 winter-elevated and 609 winter-reduced proteins (proteins with unknown BP term were excluded). Dot size represents gene count per BP category, while dot color indicates the gene ratio (proportion of proteins annotated to each BP category). Only the top significantly enriched BP terms (lowest *P*-values) are displayed for clarity. **(b)** Mosaic plot of BP enriched among winter-elevated DAPs across five PACMAD grass species. The plot summarizes the distribution of functionally annotated proteins (n = 483 of 630; unannotated proteins were excluded) across 139 biological process (BP) terms. These terms were consolidated into broader functional categories to aid interpretability. Each bar shows the relative abundance of functional categories within a species, highlighting both conserved and species-specific patterns. A Chi-square test for independence ($\chi^2[70]=97.5$, $P=0.017$) was performed to evaluate variation across species. Category labels are shown in the legend.

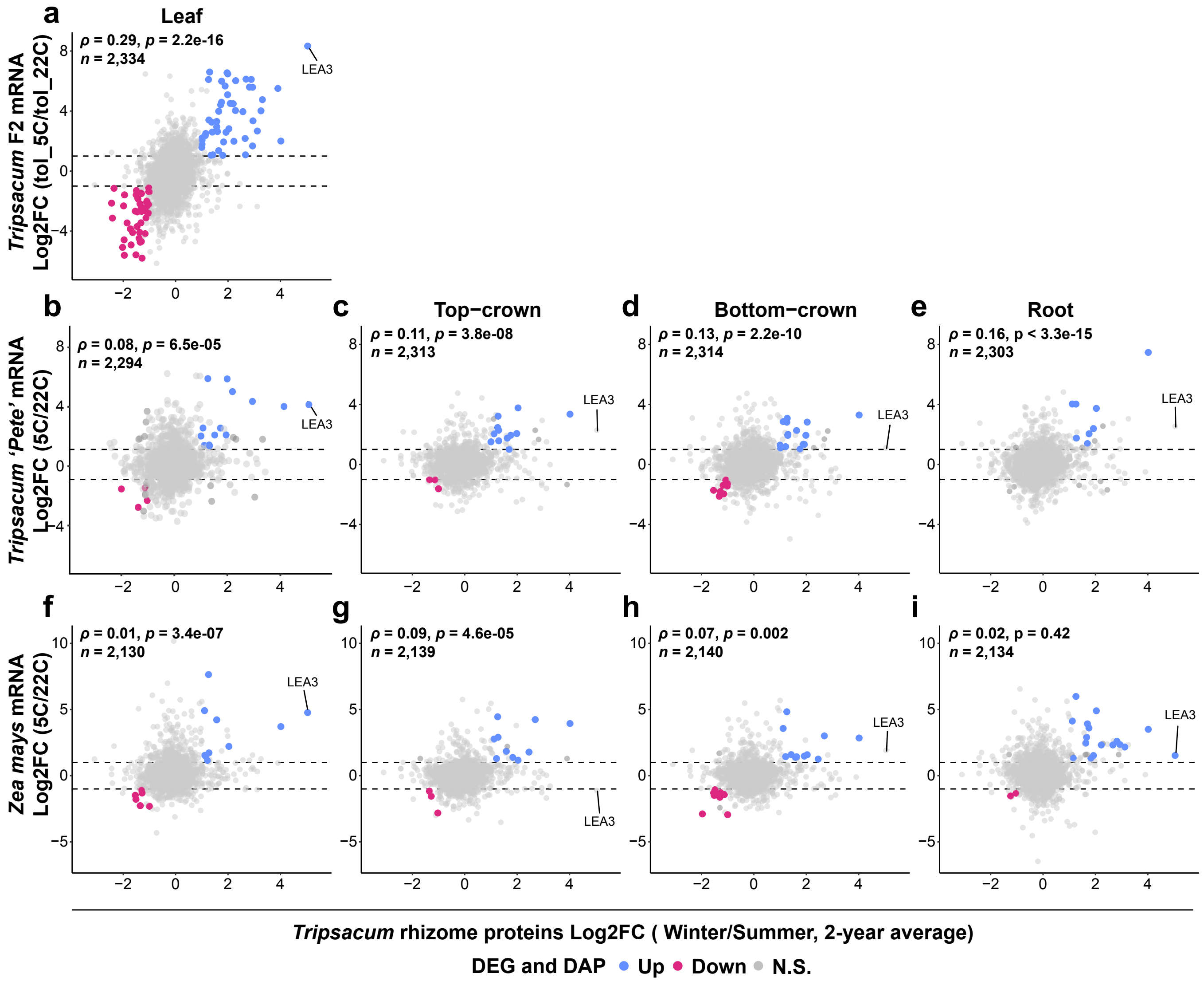


**Fig. S7. Relationship between seasonal protein abundance in rhizomes of *Tripsacum dactyloides* and *T. floridanum* hybrids and cold-responsive RNA expression in *Tripsacum* hybrids, *Tripsacum dactyloides*, and *Zea mays*.** **(a)** Scatterplots showing the relationship between *Tripsacum* rhizome proteins (x-axis, log2FC Winter/Summer, 2-year average) and *Tripsacum* leaf mRNA from a freezing tolerant $F_2$ bulk before and after 7 days cold acclimation (tol_5°C/tol_22°C) descended from the same *Tripsacum* hybrids used for rhizome proteomic analysis. **(b–e)** *Tripsacum dactyloides* (cv. 'Pete') tissues (5°C/22°C): leaf, top-crown, bottom-crown, and root. **(f–i)** *Zea mays* tissues (5°C/22°C): leaf, top-crown, bottom-crown, and root. Points are colored by shared differential expression status: blue for upregulated, pink for downregulated, and gray for non-significant changes. Dashed horizontal lines indicate the ±1 log2FC thresholds. Spearman's correlation coefficient ($\rho$) and *P*-value are shown in the upper left of each panel, along with the number of genes analyzed (*n*). The LEA3 gene is labeled in each panel.

```
                1                                                                              80
T. dactyloides  MASHQDKASYQAGETKARTEEKTGQAVGATKDTAQHAKDRASDAAGHA--------EATKQRAAETA-------EAT---
Z. mays         MASHQDKASYQAGETKARTEEKTGQAVGATKDTAQHAKDRAADAAGHAAGKGQDAKEATKQKASDTGSYLGKKTDEA---
A. gerardi      MASHQDKASYQAGETKARTEEKTGQAMGATKDTAQH------------------AKETTKQKASDTSSYLGQKTEEAKQK
S. nutans       MASHQDKASYQAGETKARTEEKTGQAMGATKDTAEH------------------AKETTKQKASDTGSYLGQKTEEA---
M. giganteus    MASHQDKASYQAGETKARTEEKTGQAMGATKDTAQH------------------AKETTKQKASDTGSYLGQKTEEA---
P. virgatum     MASHQDKASYQAGETKARTEEKAGQAMGATKDTAQHAKDRASDAAGHAAGKGHDAKEATKQKASDTGSYLGQKTDEA---

                81                                                                            160
T. dactyloides  --------KQRAAETA----------------------EATKQRAAETAEATKQKAETAEATKQKAAE----YAKDTAVS
Z. mays         --------KHKAGETT----------------------EATKQKAGETTEAAKQKAADAMEAAKQKAAEAGQYAKDTAVS
A. gerardi      AGQTTEATKQKTGETTEATKQKAGQTTEATKQKTGQTTEATKQKTGETTEAAKQKAAEAMEATKQKAAEAGQYAKETVDS
S. nutans       --------KHKTGETTEATKQKAGETT-----------EAAKQKTAETTEAAKQKTAETTEAAKQKAAEATEAAKETAVS
M. giganteus    --------KQKAGETT----------------------EAAKQKAGQTTEAAKQKAAETTEAAKQKAAEPTE--------
P. virgatum     -------TKHKAGETTEATKHKAGETTEA---------EATKQKAGETTEAAKQKTAEATKAAKQKAAEAGEYAKESAVA

                161                                                       218
T. dactyloides  GKDKSGGVIQQATEQVKSAAAGAKDAVMNTLGMGGDN-KQGDTDTT--KDSSTITRDH
Z. mays         GKDKSGGVIQQATEQVKSAAAGAKDAVMSTLGMGGDD-KQGDANTN--KDSSTITRDH
A. gerardi      GKDKSGSVIQQATEQVKSAAAGAKDAVMNTLGMGGDNNQQSDTNTN--KDSSTITRDH
S. nutans       GKDKSGGVIQQATEQVKSAAVGAKDAVMNTLGMGGDNNNQQQSDTNH-KDSSTITRDH
M. giganteus    --EKSGGVIQQATEQVKSAAAGAKDAVMNTLGMGGDNSKQGDTNTNNSKDSSTITRDH
P. virgatum     GKDKTGSAIQQATEQVKSAAVGAKDAVMSTLGMSGDN-KEGGAGNGKEEDHSTITRDQ
```

Tandem 11-mer repeat    Variant repeat (Z. mays repeat 3)

**Fig. S8. Multiple sequence alignment of LEA3 orthologs across six PACMAD species.** MAFFT alignment of LEA3 (OG0022470) protein sequences from *Tripsacum dactyloides* (Td00001aa022594), *Zea mays* (Zm00001eb294480), *Andropogon gerardi* (Ag00001aa069574), *Sorghastrum nutans* (Sn00001aa056018), *Miscanthus giganteus* (Misin17G219700), and *Panicum virgatum* (PvWBCH1.3NG170900). Blue shading indicates tandem 11-mer repeat units; orange shading indicates the variant third repeat of *Z. mays*.

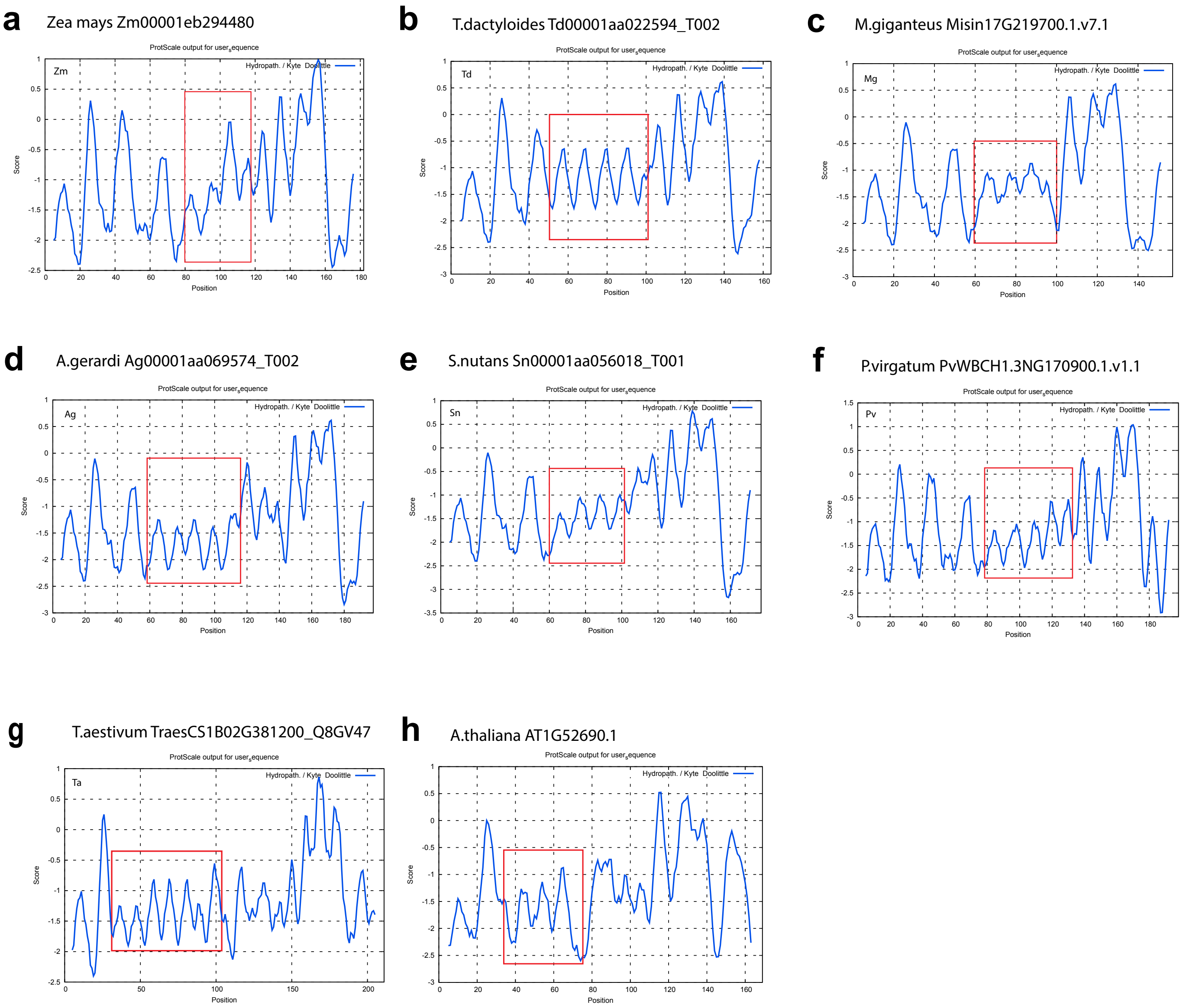


**Fig. S9**. **Hydropathy profiles of LEA3 orthologs. (a)** *Zea mays,* **(b)** *Tripsacum dactyloides*, **(c)** *Miscanthus × giganteus*, **(d)** *Andropogon gerardi*, **(e)** *Sorghastrum nutans*, **(f)** *Panicum virgatum*, **(g)** *Triticum aestivum*, and **(h)** *Arabidopsis thaliana*. Kyte-Doolittle plots were generated using ProtScale (ExPASy) with a window size of 11 amino acids. The x-axis indicates amino acid position, and the y-axis represents hydropathy score. Red boxes indicate the tandem 11-mer repeat region for each species.

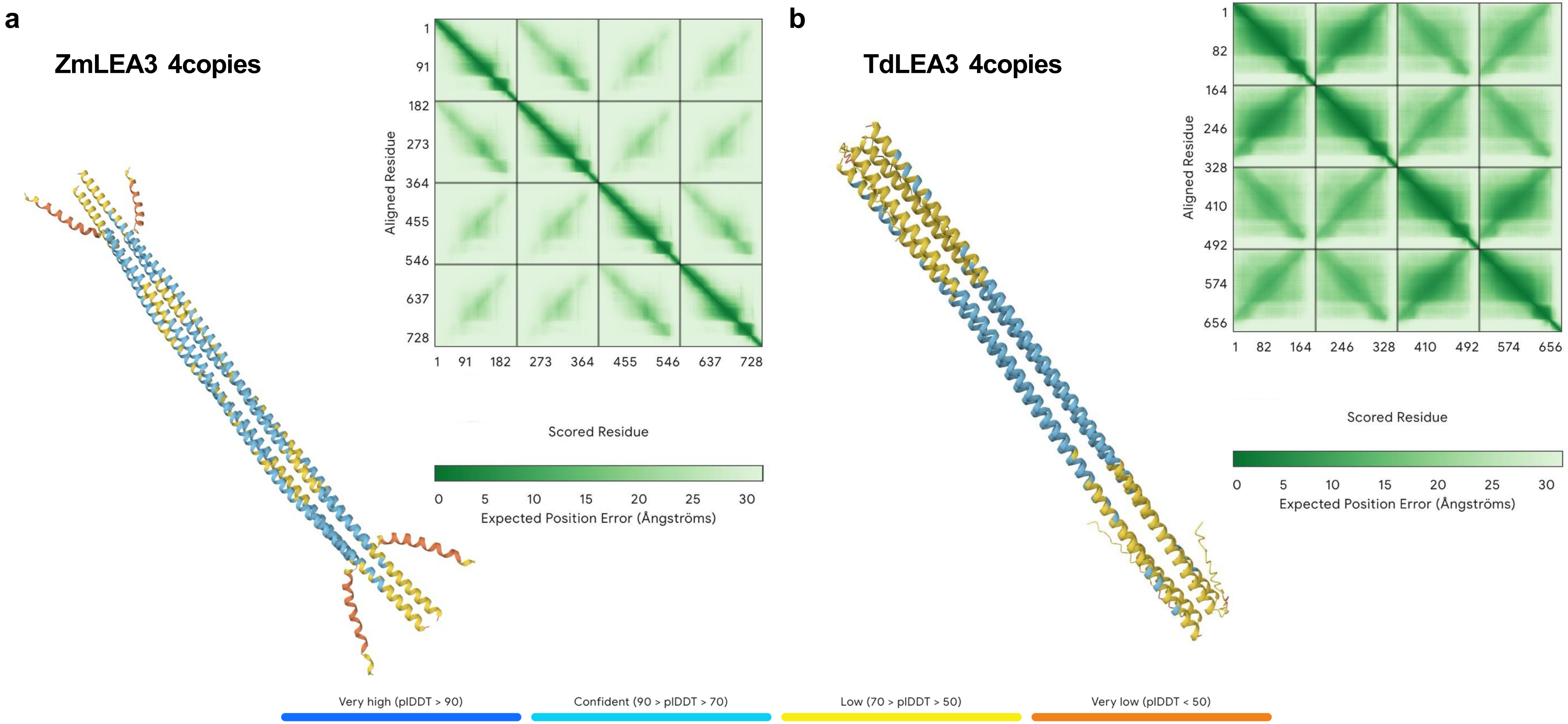


**Fig. S10. AlphaFold 3 prediction of LEA3 homo-tetramer assemblies**. Predicted quaternary structures of LEA3 homotetramers for **(a)** *Zea mays* and **(b)** *Tripsacum dactyloides*. Both orthologs are predicted to form tetrameric assemblies. Colors indicate per-residue confidence (pLDDT), with dark blue/blue representing high to very high confidence. The *Z. mays* complex displays a distinct oligomeric packing arrangement relative to *Tripsacum*. Predicted Aligned Error (PAE) plots. Low error values (dark green) in the off-diagonal regions confirm high confidence in the relative positioning of the subunits, validating the predicted tetrameric interfaces for both species.
