## Supplemental Codes for "Constrained evolution of a core winter proteome across independently cold-adapted PACMAD grasses": S1_import_Ag_Mg_data.R

```
# Script Title: Import and Split Andropogon/Miscanthus (Ag/Mg) 2022 TMT Proteomics Data
# 
# Description: Imports TMT proteomics data from Excel for Andropogon and Miscanthus
#              2022 samples, cleans column names, maps TMT labels to sample names,
#              and splits the data into separate Andropogon (Ag) and Miscanthus (Mg)
#              datasets with standardized column names.
#
# Author: Elad Oren
# Created: 2025-01-01
# Last Modified: 2025-04-01
#
# Input: 
#   - CintaRomay10474963_Ag_Set1_Fr-2_with2peptides_with-adj-p-values.xlsx
#   - Ag_Mg_tmt_labels.txt - Sample metadata and TMT label mapping
#
# Output:
#   - 0_Ag_clean-data.csv - Andropogon dataset with full data
#   - 0_Ag_clean-data_removed-NA.csv - Filtered Andropogon dataset without missing p-values
#   - 0_Mg_0_split_data_from_Ag.csv - Miscanthus dataset split from original file
#
# Dependencies: readxl, stringr, dplyr, readr
#
# Notes: The script processes a single Excel file containing both Andropogon and 
#        Miscanthus samples, separates them into species-specific datasets, and
#        standardizes statistical column names. Calculates protein detection
#        statistics for Andropogon samples by season.

setwd("~/Library/CloudStorage/OneDrive-Personal/Projects/BucklerLab/Tripsacum_freezing_tolerance/manuscript_V2/0_raw_data/Ag_Mg_2022/")

# Simplified script to import TMT proteomics data, relabel, and extract essential columns
library(readxl)    # For reading Excel files
library(stringr)   # For string manipulation
library(dplyr)     # For data manipulation
library(readr)

rm(list=ls())

# Import the protein data from Excel
protein_data <- read_excel("CintaRomay10474963_Ag_Set1_Fr-2_with2peptides_with-adj-p-values.xlsx", 
                           sheet = "Proteins")

## Clean column names
# Inspect the column names
original_colnames <- colnames(protein_data)
print("Original Column Names:")
print(original_colnames)

# Function to clean column names
clean_colnames <- function(colnames) {
  # Remove ' n/a'
  colnames <- gsub(" n/a", "", colnames)
  # Replace spaces and special characters with underscores
  colnames <- gsub("[[:space:][:punct:]]+", "_", colnames)
  # Remove multiple consecutive underscores
  colnames <- gsub("_+", "_", colnames)
  # Remove leading and trailing underscores
  colnames <- gsub("^_|_$", "", colnames)
  return(colnames)
}

# Apply the cleaning function to the column names
cleaned_colnames <- clean_colnames(original_colnames)
colnames(protein_data) <- cleaned_colnames

# Verify the column name changes
print(cleaned_colnames)


# Load your key
key <- read.delim("Ag_Mg_tmt_labels.txt", stringsAsFactors = FALSE)

# Get current colnames
all_cols <- colnames(protein_data)

# Match abundance-related columns with F1_<tmt> and season
pattern <- "^(.*)_F1_(\\d+[A-Z]?)_Sample_.*_(Autumn|Fall|Winter|Summer)$"
matched <- str_match(all_cols, pattern)

# Only keep rows with a match
col_info <- data.frame(
  old_name = all_cols[!is.na(matched[, 1])],
  prefix = matched[!is.na(matched[, 1]), 2],
  tmt_label = matched[!is.na(matched[, 1]), 3],
  season = matched[!is.na(matched[, 1]), 4],
  stringsAsFactors = FALSE
)

# Normalize "Autumn" to "Fall" for matching
col_info$season[col_info$season == "Autumn"] <- "Fall"

# Join with key to get sample_name (not botanical_accession!)
col_info <- left_join(col_info, key, by = c("tmt_label", "season"))

# Check for missing matches
if (any(is.na(col_info$sample_name))) {
  warning("Some columns did not match any entry in the key file.")
  print(col_info[is.na(col_info$sample_name), ])
}

# Build new names using sample_name
col_info <- col_info %>%
  mutate(new_name = paste0(prefix, "_", sample_name, "_", season))

# Apply renaming
rename_vector <- setNames(col_info$new_name, col_info$old_name)
colnames(protein_data)[colnames(protein_data) %in% names(rename_vector)] <-
  rename_vector[colnames(protein_data)[colnames(protein_data) %in% names(rename_vector)]]

colnames(protein_data)

#rename_protein_id
protein_data <- protein_data %>%
  rename(protein_id = Accession)

# Step 2: Drop columns with _Fall or _Autumn suffix
protein_data <- protein_data %>%
  select(-matches("(_Fall|_Autumn)$"))

# Step 3: Split into Miscan/Mg and Andro/Ag tables
# Metadata columns to keep
meta_cols <- c(
  "protein_id", "Checked", "Master", "Description", "Coverage",
  "Peptides", "PSMs", "AAs", "Modifications"
)

# Split: Miscan (Mg + Miscan stats)
miscan_protein_data <- protein_data %>%
  select(
    all_of(meta_cols),
    matches("_Mg-\\d+_"),
    matches("Miscan")
  )

# Split: Andro (Ag + Andro stats)
andro_protein_data <- protein_data %>%
  select(
    all_of(meta_cols),
    matches("_Ag-\\d+_"),
    matches("Andro")
  )


# Step 5: Clean Andro stat column names
colnames(andro_protein_data) <- colnames(andro_protein_data) %>%
  str_replace_all("Abundance_Ratio_P_Value_Andro_([^_]+)_Andro_", "pval_\\1_") %>%
  str_replace_all("Abundance_Ratio_Adj_P_Value_Andro_([^_]+)_Andro_", "adj_pval_\\1_") %>%
  str_replace_all("Abundance_Ratio_Andro_([^_]+)_Andro_", "Abundance_Ratio_\\1_") %>%
  str_replace_all("Abundances_Grouped_CV_Andro_", "Abundance_Grouped_CV_") %>%
  str_replace_all("Abundances_Grouped_Andro_", "Abundance_Grouped_")

andro_protein_data <- andro_protein_data %>%
  # Step 1: remove columns containing "Autumn"
  select(-contains("Autumn")) %>%
  
  # Step 2: reorder columns — stats first, then samples
  select(
    protein_id, Checked, Master, Description, Coverage, Peptides, PSMs, AAs, Modifications,
    matches("^Abundance_Ratio_"),
    matches("^pval_"),
    matches("^adj_pval_"),
    matches("^Abundance_Grouped"),
    everything()  # puts all remaining columns (samples etc.) after
  )


# Save the original dataset with renamed columns
write.csv(andro_protein_data, "0_Ag_clean-data.csv", row.names = FALSE)

#####statisctics for proptein counts

# Step 1: Select protein_id + Abundances_Normalized columns
df_norm <- andro_protein_data %>%
  select(protein_id, starts_with("Abundances_Normalized_"))

# Step 2: Count proteins > 0 per sample
counts_per_sample <- df_norm %>%
  select(-protein_id) %>%
  summarise(across(
    everything(),
    ~ sum(.x > 0, na.rm = TRUE)
  )) %>%
  pivot_longer(
    cols = everything(),
    names_to = "sample",
    values_to = "num_detected_proteins"
  )

# Step 3: Extract species and season
counts_per_sample <- counts_per_sample %>%
  mutate(
    species = str_extract(sample, "Ag|Mg"),  # Extract species (add more if needed)
    season = str_extract(sample, "Summer|Winter")        # Extract season
  )

# Step 4: Summarize by species + season
summary_per_species_season <- counts_per_sample %>%
  group_by(species, season) %>%
  summarise(
    mean_detected = mean(num_detected_proteins),
    sd_detected = sd(num_detected_proteins),
    .groups = "drop"
  )

######

# Step 3: Drop rows with any NA in adj_pval columns
andro_protein_data <- andro_protein_data %>%
filter(if_all(starts_with("adj_pval_"), ~ !is.na(.)))


# Save the original dataset with renamed columns

write.csv(andro_protein_data, "0_Ag_clean-data_removed-NA.csv", row.names = FALSE)


# Step 4: Clean Miscan stat column names
colnames(miscan_protein_data) <- colnames(miscan_protein_data) %>%
  str_replace_all("Abundance_Ratio_P_Value_Miscan_([^_]+)_Miscan_", "pval_\\1_") %>%
  str_replace_all("Abundance_Ratio_Adj_P_Value_Miscan_([^_]+)_Miscan_", "adj_pval_\\1_") %>%
  str_replace_all("Abundance_Ratio_Miscan_([^_]+)_Miscan_", "Abundance_Ratio_\\1_") %>%
  str_replace_all("Abundances_Grouped_CV_Miscan_", "Abundance_Grouped_CV_") %>%
  str_replace_all("Abundances_Grouped_Miscan_", "Abundance_Grouped_")

miscan_protein_data <- miscan_protein_data %>%
  # Step 1: Remove Autumn-related columns
  select(-contains("Autumn")) %>%
  
  # Step 2: Reorder columns — stats first, then samples
  select(
    protein_id, Checked, Master, Description, Coverage, Peptides, PSMs, AAs, Modifications,
    matches("^Abundance_Ratio_"),
    matches("^pval_"),
    matches("^adj_pval_"),
    matches("^Abundance_Grouped"),
    everything()
  ) #%>%
  
  # Step 3: Drop rows with any NA in adj_pval columns
  #filter(if_all(starts_with("adj_pval_"), ~ !is.na(.)))

# Save the original dataset with renamed columns
write.csv(miscan_protein_data, "0_Mg_0_split_data_from_Ag.csv", row.names = FALSE)
```
