## Supplemental Codes for "Constrained evolution of a core winter proteome across independently cold-adapted PACMAD grasses": S2_map_Mg_peptides_to_proteins.R

```
# Script Title: Map Miscanthus (Mg) Peptides to Andropogon (Ag) Proteins
# 
# Description: Maps peptide-level BLAST hits to identify the best Miscanthus 
#              protein match for each Andropogon protein using a multi-level 
#              tie-breaking approach. Prioritizes matches by: 1) number of 
#              distinct peptides, 2) sum of bitscores, and 3) median 
#              percent identity.
#
# Author: Elad Oren
# Created: 2025-01-01
# Last Modified: 2025-04-01
#
# Input: 
#   - Ag_Set1_peptides_vs_Mg_results_ultrasensitive.tsv - DIAMOND BLAST results of 
#     Andropogon peptides against Miscanthus proteins
#
# Output:
#   - 0_Mg_1_best_mapping_peptides.csv - Best Miscanthus protein match for 
#     each Andropogon protein
#
# Dependencies: readr, dplyr
#
# Notes: This script performs a deterministic selection of the best Mg protein
#        for each Ag protein based on three sequential criteria. The results
#        serve as a mapping table for downstream cross-species comparisons.

setwd("~/Library/CloudStorage/OneDrive-Personal/Projects/BucklerLab/Tripsacum_freezing_tolerance/manuscript_V2/0_raw_data/Ag_Mg_2022/")

rm(list=ls())


# 1) Load libraries
library(readr)   # for read_tsv()
library(dplyr)   # for data manipulation (group_by, summarize, etc.)


# 2) Read the TSV file (adjust file path as needed)
file_path <- "Ag_Set1_peptides_vs_Mg_results_ultrasensitive.tsv"
df <- read_tsv(file_path)  # make sure your columns match expected headers

# Quick sanity check
cat("\nInitial data frame:\n")
print(dim(df))        # rows x cols
print(head(df, 5))    # first few rows

# 3) Aggregate at (Ag_protein, Mg_protein_id)
#    - Count distinct peptides
#    - Sum bitscores
#    - Compute median pident
df_rollup <- df %>%
  group_by(Ag_protein, Mg_protein_id) %>%
  summarize(
    n_peptides = n_distinct(Ag_peptides),
    sum_bitscore = sum(bitscore),
    median_pident = median(pident),
    .groups = "drop"        # finalize the grouping
  )

cat("\nRolled-up data (first 10 rows):\n")
print(head(df_rollup, 10))

# 4) Sort within each Ag_protein by:
#    1) n_peptides (descending),
#    2) sum_bitscore (descending),
#    3) median_pident (descending)
# Then pick the top row from each group to get the single best Mg match.
df_best <- df_rollup %>%
  group_by(Ag_protein) %>%
  arrange(
    desc(n_peptides),
    desc(sum_bitscore),
    desc(median_pident),
    .by_group = TRUE
  ) %>%
  slice_head(n = 1) %>%
  ungroup()

cat("\nBest Mg protein for each Ag_protein (first 10 rows):\n")
print(head(df_best, 10))


write_csv(df_best, "0_Mg_1_best_mapping_peptides.csv")
```
