## Supplemental Codes for "Constrained evolution of a core winter proteome across independently cold-adapted PACMAD grasses": S3_summarize_Mg_proteins.R

```
# Script Title: Summarize Miscanthus (Mg) Protein Data Based on Peptide Mapping
# 
# Description: Consolidates Miscanthus proteomics data by combining the split
#              Miscanthus dataset with peptide mapping information. For each
#              Miscanthus protein, summarizes peptide coverage metrics and
#              aggregates abundance values from multiple Andropogon proteins
#              that map to the same Miscanthus protein.
#
# Author: Elad Oren
# Created: 2025-01-01
# Last Modified: 2025-04-01
#
# Input: 
#   - 0_Mg_0_split_data_from_Ag.csv - Extracted Miscanthus data
#   - 0_Mg_1_best_mapping_peptides.csv - Peptide mapping between Ag and Mg
#
# Output:
#   - 0_Mg_2_clean-data.csv - Summarized Miscanthus protein data
#   - 0_Mg_2_clean-data_removed-NA.csv - Filtered dataset without infinite p-values
#
# Dependencies: readr, dplyr
#
# Notes: The script aggregates data by Miscanthus protein ID, summing abundances
#        and selecting the minimum p-values. Calculates protein detection
#        statistics by species and season, and removes entries with infinite
#        adjusted p-values.


library(readr)
library(dplyr)

rm(list=ls())


# ------------------------------
# 1. Read the main "clean" Mg data
# ------------------------------
df_mg <- read_csv("0_Mg_0_split_data_from_Ag.csv")

df_mg <- df_mg %>%
  rename(Ag_protein = protein_id)

# Quick check
cat("df_mg columns:\n")
print(colnames(df_mg))
cat("\nFirst few rows:\n")
print(head(df_mg))

# ------------------------------
# 2. (Optional) Read a second table with Mg_protein_id (if needed)
#    and JOIN them. If your df_mg file already has all the columns
#    you need, you can skip this step.
# ------------------------------
# Suppose you have df_best with columns: Mg_protein_id, something else
df_best <- read_csv("0_Mg_1_best_mapping_peptides.csv")
 
df_joined <- df_mg %>%
   left_join(df_best, by = "Ag_protein")

# If you don't actually need a separate join, just proceed with df_mg:
#df_joined <- df_mg

# ------------------------------
# 3. Summarize by Mg_protein_id
#    We'll assume your CSV has columns named, for example:
#    "pval"            -> p-value for each row
#    "ratio"           -> ratio for each row
#    "Abundances (Grouped): Miscan_Winter"
#    "Abundances (Grouped): Miscan_Summer"
# ------------------------------


df_joined <- df_joined %>%
  filter(!is.na(Mg_protein_id))  # Remove rows with NA in Mg_protein_id

df_summary <- df_joined %>%
  group_by(Mg_protein_id) %>%
  summarize(
    Peptides = sum(n_peptides, na.rm = TRUE),
    Coverage = median(median_pident, na.rm = TRUE),
    Bitscore = sum(sum_bitscore, na.rm = TRUE),
    
    # average the ratio column
    Abundance_Ratio_Winter_Summer = sum(Abundance_Grouped_Winter) / sum(Abundance_Grouped_Summer),

    # pick the minimum p-value across all rows for that Mg protein
    pval = min(pval_Winter_Summer, na.rm = TRUE),
    padj = min(adj_pval_Winter_Summer, na.rm = TRUE),
    
    # sum up the grouped winter and summer abundances
    Abundance_Grouped_Winter = sum(Abundance_Grouped_Summer, na.rm = TRUE),
    Abundance_Grouped_Winter = sum(Abundance_Grouped_Winter, na.rm = TRUE),
    `Abundance_Mg-1_Summer` = sum(`Abundance_Mg-1_Summer`, na.rm = TRUE),
    `Abundance_Mg-2_Summer` = sum(`Abundance_Mg-2_Summer`, na.rm = TRUE),
    `Abundance_Mg-3_Summer` = sum(`Abundance_Mg-3_Summer`, na.rm = TRUE),
    `Abundance_Mg-1_Winter` = sum(`Abundance_Mg-1_Winter`, na.rm = TRUE),
    `Abundance_Mg-2_Winter` = sum(`Abundance_Mg-2_Winter`, na.rm = TRUE),
    `Abundance_Mg-3_Winter` = sum(`Abundance_Mg-3_Winter`, na.rm = TRUE),
    `Abundances_Normalized_Mg-1_Summer` = sum(`Abundances_Normalized_Mg-1_Summer`, na.rm = TRUE),
    `Abundances_Normalized_Mg-2_Summer` = sum(`Abundances_Normalized_Mg-2_Summer`, na.rm = TRUE),
    `Abundances_Normalized_Mg-3_Summer` = sum(`Abundances_Normalized_Mg-3_Summer`, na.rm = TRUE),
    `Abundances_Normalized_Mg-1_Winter` = sum(`Abundances_Normalized_Mg-1_Winter`, na.rm = TRUE),
    `Abundances_Normalized_Mg-2_Winter` = sum(`Abundances_Normalized_Mg-2_Winter`, na.rm = TRUE),
    `Abundances_Normalized_Mg-3_Winter` = sum(`Abundances_Normalized_Mg-3_Winter`, na.rm = TRUE),
    
    
    .groups = "drop"
  ) 

df_summary <- df_summary %>%
  rename(protein_id = Mg_protein_id)

df_summary$Abundance_Ratio_Winter_Summer <- df_summary$Abundance_Grouped_Winter / df_summary$Abundance_Grouped_Summer


#write_csv(df_joined, "df_joined_Mg_Ag_peptides.csv")
write_csv(df_summary, "0_Mg_2_clean-data.csv")

#####statisctics for proptein counts

# Step 1: Select protein_id + Abundances_Normalized columns
df_norm <- df_summary %>%
  select(protein_id, starts_with("Abundances_Normalized_"))

# Step 2: Count proteins > 0 per sample
counts_per_sample <- df_norm %>%
  select(-protein_id) %>%
  summarise(across(
    everything(),
    ~ sum(.x > 0, na.rm = TRUE)
  )) %>%
  pivot_longer(
    cols = everything(),
    names_to = "sample",
    values_to = "num_detected_proteins"
  )

# Step 3: Extract species and season
counts_per_sample <- counts_per_sample %>%
  mutate(
    species = str_extract(sample, "Ag|Mg"),  # Extract species (add more if needed)
    season = str_extract(sample, "Summer|Winter")        # Extract season
  )

# Step 4: Summarize by species + season
summary_per_species_season <- counts_per_sample %>%
  group_by(species, season) %>%
  summarise(
    mean_detected = mean(num_detected_proteins),
    sd_detected = sd(num_detected_proteins),
    .groups = "drop"
  )

######


# Step 3: Drop rows with any NA in adj_pval columns
df_summary <- df_summary %>%
  filter(if_all(starts_with("padj"), ~ !is.infinite(.)))


write_csv(df_summary, "0_Mg_2_clean-data_removed-NA.csv")
```
